# *Salmonella* Effector SteE Reprogrammes the Macrophage Regulatory Network to Drive Specific Hyperactivation of STAT3 Target Genes

**DOI:** 10.1101/2025.07.11.664171

**Authors:** Ines Diaz-del-Olmo, Paul A. O’Sullivan, Sayaka Shizukuishi, Adriana Stypulkowska, Ioanna Panagi, Krzysztof Grzymajlo, Michinaga Ogawa, Peter W. S. Hill, Teresa L. M. Thurston

**Affiliations:** Sir William Dunn School of Pathology, University of Oxford, South Parks Road, Oxford OX1 3RE, UK; Department of Infectious Disease, Centre for Bacterial Resistance Biology, Imperial College London, London, SW7 2AZ, UK; Bacterial Pathogenesis and Immune Signalling Laboratory, The Francis Crick Institute, 1 Midland Road, London, NW1 1AT, UK; Department of Bacteriology, National Institute of Infectious Diseases, Tokyo, Japan; Wroclaw University of Environmental and Life Sciences, Faculty of Veterinary Medicine, Department of Biochemistry and Molecular Biology, Wrocław, Poland; Department of Infectious Diseases, School of Immunology and Microbial Sciences, Guy’s Hospital, King’s College London, London, SE1 9RT, UK

**Keywords:** host-pathogen interactions, *Salmonella*, macrophage, anti-inflammatory, STAT3, virulence factor, effector, SteE, gene regulatory network

## Abstract

The ability of *Salmonella* Typhimurium to exploit macrophages as a niche for survival, replication and dissemination is central to its pathogenesis. The effector SteE, which polarises macrophages into an anti-inflammatory state, is critical during invasive disease. SteE operates via an unprecedented mechanism, reprogramming the host serine/threonine kinase GSK3 to perform tyrosyl-directed phosphorylation of neosubstrates, including the immune transcription factors STAT1 and STAT3. Here, we demonstrate that SteE-driven transcriptional reprogramming relies critically and specifically on STAT3 phosphorylation and DNA binding. By activating STAT3 via a non-canonical pathway, bypassing endogenous negative feedback mechanisms, SteE drives hyperactivation of STAT3 target genes, surpassing the effects of canonical IL10 signalling. Hyperactivation correlates with elevated phosphorylated STAT3 in the macrophage nucleus, facilitating opening of chromatin regions not accessible during endogenous cytokine signalling. Overall, our study illustrates how hijacking of a signalling pathway by SteE dramatically reshapes the macrophage gene regulatory network to enhance *Salmonella* immune evasion.

## INTRODUCTION

*Salmonella* Typhimurium, a Gram-negative facultative intracellular pathogen, causes millions of non-typhoidal *Salmonella* (NTS) infections globally each year. Whereas most cases remain confined to the gastrointestinal tract, a significant subset progress to invasive non-typhoidal *Salmonella* (iNTS) disease, particularly in vulnerable groups such as young children and individuals with primary immunodeficiencies or advanced HIV infection (Stanaway et al., 2019). iNTS disease is characterised by systemic bacterial dissemination to organs such as the spleen, liver, and bone marrow, and has a high mortality rate of 15– 20%, resulting in over 70,000 deaths annually (Stanaway et al., 2019). This burden is particularly acute in sub-Saharan Africa, where iNTS is a leading cause of bloodstream infections (Stanaway et al., 2019).

The ability of *Salmonella* to exploit macrophages as niches for survival, replication, and dissemination is central to iNTS pathogenesis (Fields et al., 1986; Richter-Dahlfors et al., 1997; Salcedo et al., 2001; Monack, 2013). This process relies on the *Salmonella* Pathogenicity Island 2 (SPI2)-encoded type 3 secretion system (T3SS), which delivers effector proteins into host cells to reprogramme their functions (Jennings et al., 2017). Among these effectors, SteE plays a key role in polarising macrophages into an anti-inflammatory, M2-like state (Jaslow et al., 2018; Panagi et al., 2020; Pham et al., 2020, 2023; Stapels et al., 2018). Animal studies revealed that SteE-induced macrophage reprogramming enhances bacterial persistence and dissemination by counteracting local pro-inflammatory signals in the macrophage microenvironment which, otherwise, would drive bacterial clearance, such as Tumour Necrosis Factor α (TNF-α) and type I interferons (Pham et al., 2020, 2023). The absence of SteE impairs both acute and persistent systemic infection, underscoring its pivotal role in invasive *Salmonella* Typhimurium pathogenesis (Gibbs et al., 2020; Pham et al., 2020). In addition, the activity of SteE is instrumental during antibiotic treatment, where reprogrammed macrophages act as a niche for *Salmonella*, enabling bacterial survival and relapse after therapy ends (Stapels et al., 2018). These findings suggest that SteE complicates pathogen eradication and contributes to bacterial persistence, highlighting the need for a deeper understanding of its molecular mechanisms to inform therapeutic development.

Mechanistically, SteE induces the host serine/threonine kinase GSK3 to directly phosphorylate neosubstrates on a tyrosine residue (Panagi et al., 2020). Two transcription factors involved in immune regulation, STAT1 and STAT3, traditionally associated with pro-inflammatory and anti-inflammatory roles, respectively, have been identified as phosphorylation targets following translocation of SteE (Gibbs et al., 2020; Jaslow et al., 2018; Panagi et al., 2020; Stepien et al., 2024). Emerging evidence also suggests that SteE activity may directly repress pro-inflammatory transcriptional pathways in macrophages (Heyman et al., 2023). These findings highlight the potential of SteE to modulate macrophage signalling and gene regulation through multiple interconnected mechanisms, underscoring the need for further investigation to fully understand its molecular targets and functions.

In this study, we complete the regulatory pathway linking SteE activity to host transcriptional reprogramming in macrophages. We show that SteE translocation results in the opening of chromatin regions associated with anti-inflammatory gene activation, driving M2-like macrophage responses through chromatin remodelling. We identify STAT3 as the critical and specific driver of transcriptional changes during SteE-mediated reprogramming and suggest that STAT1 phosphorylation represents a non-functional biological artefact. Most strikingly, we show that SteE-induced STAT3 phosphorylation and DNA binding leads to hyperactivation of STAT3 target genes that surpasses the effects of canonical IL10 signalling. This hyperactivation is linked to increased phosphorylated STAT3 in the macrophage nucleus and accessibility of chromatin regions not opened by endogenous cytokine signalling. Together, our results illustrate how, by hyperactivating STAT3 via a non-canonical pathway that bypasses endogenous negative feedback loops, SteE dramatically rewires the macrophage gene regulatory network to enhance immune evasion and persistence by *Salmonella* Typhimurium.

## RESULTS

### SteE drives SPI2-dependent anti-inflammatory gene activation in macrophages

Although the role of SteE in SPI2-dependent M2-like macrophage polarisation is well-documented, recent evidence suggests its activity extends beyond this established function. For example, emerging data hint that SteE modulates both pro-inflammatory and anti-inflammatory pathways (Gibbs et al., 2020; Jaslow et al., 2018; Panagi et al., 2020; Stepien et al., 2024), raising questions about the breadth of its influence and the mechanisms it employs. This led us to examine two critical questions: (1) how extensive is the activity of SteE on macrophage phenotypes, and (2) which transcriptional mediators does SteE target to achieve its reprogramming effects?

To address the first question, we used RNA-Sequencing (RNA-seq) to examine SPI2-dependent transcriptional changes induced in mouse primary bone marrow-derived macrophages (pBMDMs) 18 hours after *Salmonella* uptake. Macrophages were infected with either wild-type *Salmonella* or an isogenic *ssaV* mutant that cannot translocate SPI2 effectors. This analysis identified over 2,000 genes whose expression was influenced by SPI2 effector translocation in macrophages infected with both the invasive ST313 (D23580) and gastrointestinal-associated ST19 (12023/14028) strains of *Salmonella* Typhimurium, with nearly equal numbers of genes being activated or repressed (Figure 1A and S1A-B, Table S1). To evaluate the role of SteE in SPI2-dependent transcriptional changes, we compared gene expression in macrophages infected with *steE*-mutant *Salmonella* to those infected with wild-type bacteria. Of the SPI2-repressed genes, only 4 showed a statistically significant (adj. p-value <0.01) and biologically meaningful (>2-fold) loss of repression in *steE*-mutant infected macrophages (Figure 1B, Table S1). In contrast, hundreds of SPI2-activated genes were dependent on SteE (Figure 1B, Table S1). Consistently, principal component analysis revealed that *steE*-mutant infected macrophages clustered with wild-type infected macrophages based on SPI2-repressed genes – but with *ssaV*-mutant infected macrophages for SPI2-activated genes (Figure 1C).

**Figure 1.**
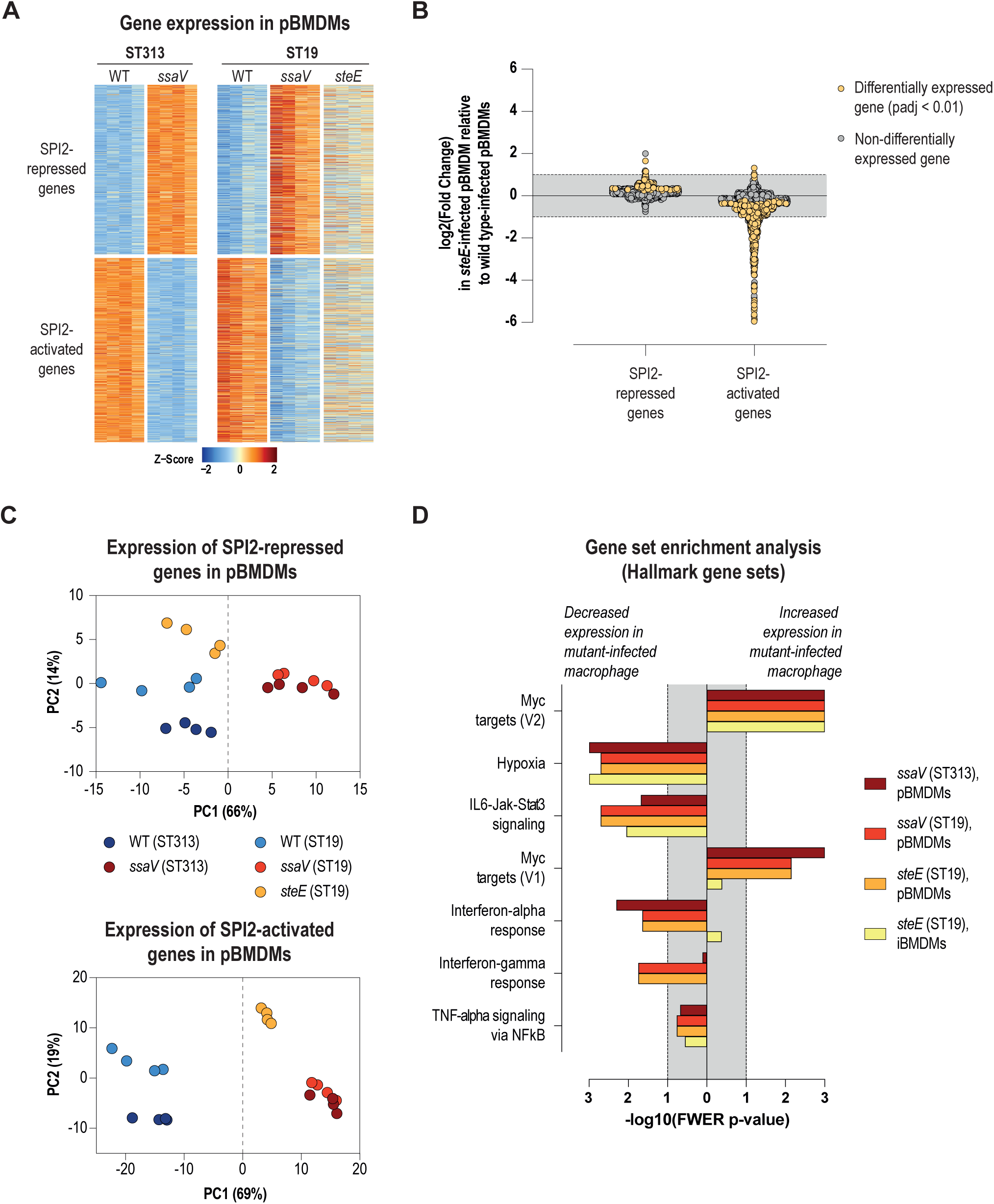
SteE drives SPI2-dependent transcriptional activation in infected macrophages. **(A-D)** Primary bone marrow-derived macrophages (pBMDMs) were infected with wild-type or *ssaV*-mutant ST313 *Salmonella* Typhimurium (D23580), or wild-type, *ssaV*-mutant or *steE*-mutant ST19 *Salmonella* Typhimurium (12023/14028). RNA sequencing was performed on samples taken 18 hours post-uptake. **(A)** Heatmap showing clusters of genes that are differentially expressed (adjusted *p*-value < 0.01, DESeq2) in pBMDMs infected with wild-type *Salmonella* Typhimurium compared to *ssaV* mutant-infected pBMDMs. Genes are classified as SPI2-activated or SPI2-repressed based on their expression patterns. Colours represent Z-scores, reflecting relative gene expression levels across all samples. **(B)** Log_2_ fold change in gene expression of SPI2-repressed and SPI2-activated genes (as identified in panel A) in pBMDMs infected with *steE*-mutant ST19 relative to wild-type ST19. Genes meeting the threshold for differential expression (adjusted *p*-value < 0.01, DESeq2) are highlighted in yellow. **(C)** Principal component analysis (PCA) of SPI2-repressed (upper panel) and SPI2-activated (lower panel) genes in pBMDMs infected with the indicated *Salmonella* Typhimurium strains. The percentage of variance explained by the first (PC1) and second (PC2) principal components is shown. **(D)** Gene set enrichment analysis (GSEA) identifying hallmark pathways with significantly increased or decreased activity (FWER-adjusted *p*-value < 0.1). Comparisons were made between pBMDMs infected with *ssaV*-mutant strains (ST19 or ST313) and their respective isogenic wild-type *Salmonella* strains, as well as between primary (pBMDMs) or immortalised (iBMDMs) macrophages infected with *steE*-mutant ST19 and its isogenic wild-type counterpart.

Gene set enrichment analysis (GSEA) indicated that *steE*-mutant-infected macrophages exhibited significantly reduced expression of hypoxia and IL6-JAK-STAT3 signalling pathway targets compared to wild-type-infected macrophages (Figure 1D Table S2). As expected, genes associated with inflammatory pathways, including TNF-α and type I interferon responses, were strongly upregulated in *Salmonella*-infected macrophages relative to naïve (unchallenged) macrophages (Figure S1C, Table S2). However, in contrast with previous suggestions (Heyman et al., 2023), we observed no consistently significant changes in type I interferon or TNF-α responses in *steE*-mutant-infected primary and immortalised BMDMs compared to those infected with an isogenic wild-type strain (Figure 1D, Table S2).

Collectively, these findings suggest that the primary function of SteE is to drive an anti-inflammatory gene expression programme in macrophages. The proposed ability of SteE to exert transcriptional activation or repression of pro-inflammatory genes likely represents a secondary consequence of the effector’s broader capacity to reshape the macrophage gene expression landscape.

### SteE-mediated anti-inflammatory transcriptional activation is coupled to chromatin remodelling

We next set out to understand how SteE translocation drives transcriptional activation in macrophages. In eukaryotes, transcriptional regulation is primarily controlled at the chromatin level, where transcription factors bind accessible promoter and enhancer regions to regulate gene expression (Lawrence & Natoli, 2011; Thurman et al., 2012; Zaret & Carroll, 2011). Most eukaryotic DNA is tightly packed in inaccessible heterochromatin, with active promoters and enhancers having undergone chromatin remodelling to become accessible (Lawrence & Natoli, 2011; Thurman et al., 2012; Zaret & Carroll, 2011). Therefore, we complemented our RNA-Seq data with matched ATAC-Seq (Assay for Transposase Accessible Chromatin Sequencing) datasets (Buenrostro et al., 2013). This analysis revealed over 2,000 chromatin regions whose accessibility was altered by SPI2 effector translocation (Figure 2A and S2A-B, Table S3). Like the gene expression changes observed, approximately equal numbers of chromatin regions became relatively closed (∼1,000) or opened (∼1,300) following SPI2 effector translocation in macrophages infected with both the invasive ST313 and gastrointestinal-associated ST19 strains of *Salmonella* Typhimurium (Figure 2A, Table S3).

**Figure 2.**
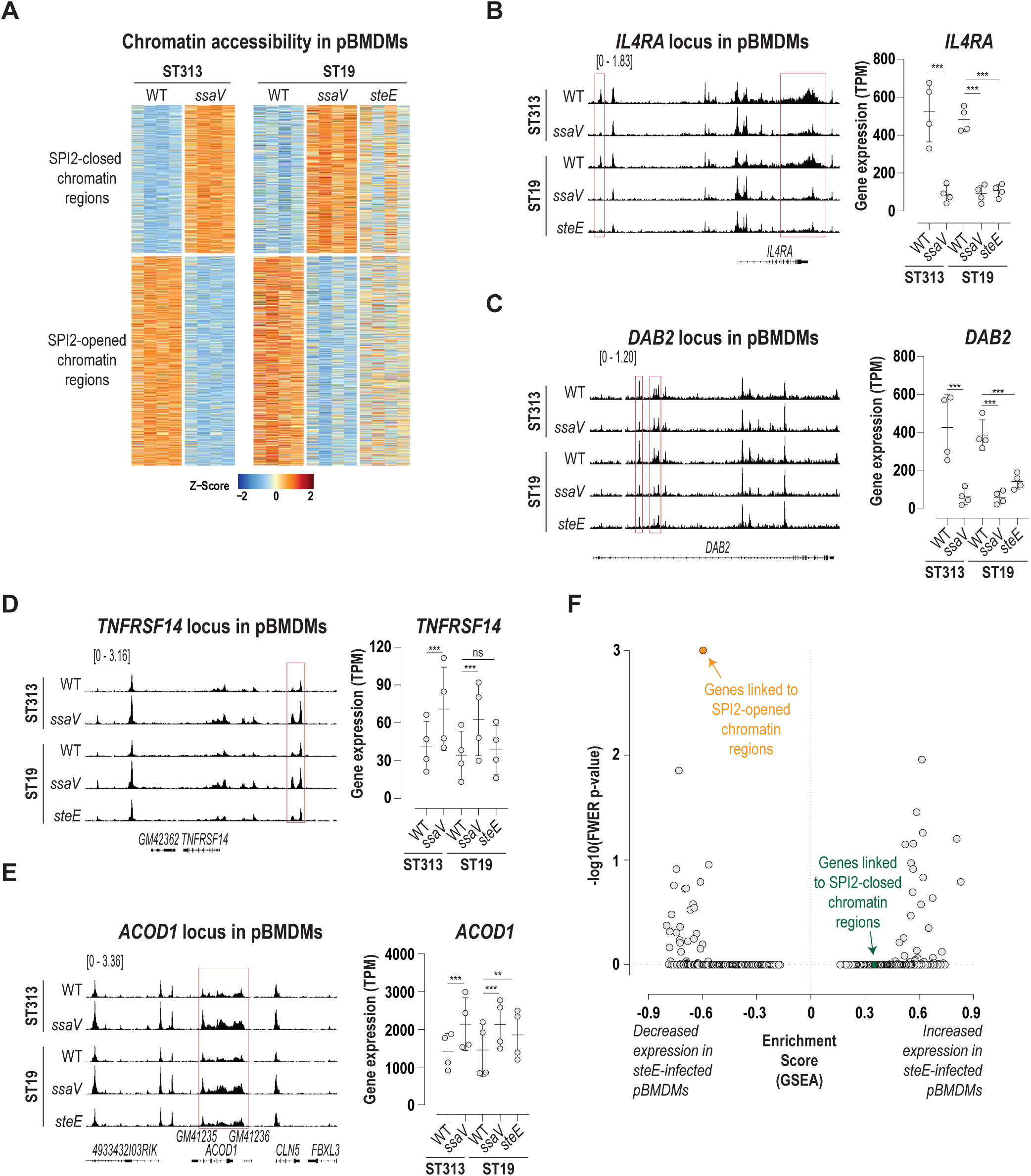
Transcriptional activation driven by SteE is coupled to chromatin remodelling. **(A-F)** Primary bone marrow-derived macrophages (pBMDMs) were infected with wild-type or *ssaV*-mutant ST313 *Salmonella* Typhimurium (D23580), or wild-type, *ssaV*-mutant or *steE*-mutant ST19 *Salmonella* Typhimurium (12023/14028). ATAC (**<u>A</u>**ssay for **<u>T</u>**ransposase **<u>A</u>**ccessible **<u>C</u>**hromatin) sequencing was performed on samples taken 18 hours post-uptake. **(A)** Heatmap showing clusters of chromatin regions that are differentially accessible (adjusted *p*-value < 0.1, DESeq2) in pBMDMs infected with wild-type *Salmonella* Typhimurium compared to *ssaV*-mutant infected pBMDMs. Regions are classified as SPI2-opened or SPI2-closed based on their accessibility patterns. Colours represent Z-scores, reflecting relative chromatin accessibility levels across all samples. **(B-E)** Local chromatin accessibility profiles (left panels; ATAC-Seq) and gene expression levels (right panels; RNA-Seq) for the anti-inflammatory marker genes *IL4RA* **(B)** and *DAB2* **(C),** as well as for the pro-inflammatory marker genes *TNFRSF14* **(D)** and *ACOD1* **(E)**, in pBMDMs infected with the indicated *Salmonella* Typhimurium strains. Red boxes highlight regions containing differentially accessible chromatin sites (DESeq2, adjusted p-value < 0.1). Differential gene expression was analysed using DESeq2, with adjusted p-values shown (∗∗∗ adj. p < 0.001, ∗∗ adj. p < 0.01, ns = not significant). **(F)** Gene set enrichment analysis (GSEA) identifying macrophage gene sets with increased or decreased expression (based on RNA-Seq; see Figure 1) in pBMDMs infected with *steE*-mutant ST19 *Salmonella* Typhimurium compared to those infected with the isogenic wild-type strain. Gene sets linked to SPI2-opened and SPI2-closed chromatin regions (see A) were identified using the Genomic Regions Enrichment of Annotations Tool (GREAT). Additional gene sets, including all hallmark gene sets, canonical pathway gene sets, and gene ontology (GO) gene sets, were sourced from GSEA. The plot shows the GSEA enrichment score (x-axis) and the -log10(FWER-adjusted *p*-value) (y-axis) for all gene sets analysed.

To assess the role of SteE in SPI2-dependent chromatin changes, we compared chromatin accessibility in pBMDMs infected with *steE*-mutant *Salmonella* to those infected with wild-type bacteria. Consistent with our RNA-Seq results, chromatin accessibility at SPI2-closed regions was only minimally affected, whereas we observed reduced chromatin accessibility at SPI2-opened regions in *steE*-mutant infected macrophages (Figures 2A-E and Figure S2D). Using the Genomic Regions Enrichment of Annotations Tool (GREAT) (McLean et al., 2010; Tanigawa et al., 2022), we linked SPI2-opened and SPI2-closed chromatin regions to their putatively associated genes (Table S4). Strikingly, of all gene sets analysed (including all hallmark, gene ontology, and canonical pathway gene sets), genes linked to SPI2-opened chromatin regions represented the genes that were most significant in terms of requiring SteE for their transcriptional upregulation (Figure 2F, Table S2). This is exemplified by the canonical anti-inflammatory genes *IL4RA* and *DAB2*, where transcriptional changes closely correspond to local chromatin accessibility alterations (Figures 2B and C). In contrast, genes associated with chromatin regions that are closed in a SPI2-dependent manner were transcriptionally upregulated in *ssaV*-mutant infected macrophages but were either unaffected or only minimally impacted by the absence of SteE (Figure 2F and S2C). This is exemplified by the pro-inflammatory genes *TNFRSF14* and *ACOD1* (Figure 2D and E). Together, these findings indicate that SteE-associated chromatin opening is functionally linked to the effector’s ability to transcriptionally activate an anti-inflammatory gene expression programme in macrophages.

### SteE-mediated transcriptional activation critically and specifically requires STAT3

Given the link between SteE-associated chromatin opening and the effector’s role in activating an anti-inflammatory gene expression programme, we next examined the features of opened chromatin regions to identify, in an unbiased manner, potential transcriptional mediators hijacked by SteE. Interestingly, although SteE reportedly phosphorylates STAT1 (Stepien et al., 2024) and activates hypoxia-associated genes (Figure 1D), SPI2-opened chromatin regions were neither enriched for STAT1 nor HIF1A binding motifs (Figure 3A, Table S5). Furthermore, footprinting analysis showed no evidence of SteE-dependent binding of these transcription factors across SPI2-regulated chromatin regions (Figure 3B, Table S6). Instead, SPI2-opened regions were strongly enriched for STAT3, STAT4, and STAT5 binding motifs, but not for motifs associated with other STAT family transcription factors (Figure 3A, Table S5). Consistently, footprinting analysis predicted a significant reduction in STAT3, STAT4, and STAT5 binding across SPI2-regulated regions in macrophages infected with *steE-*mutant *Salmonella* Typhimurium relative to a wild-type strain (Figure 3B, Table S6). Since the binding motifs for STAT3, STAT4, and STAT5 are highly similar (Kulakovskiy et al., 2018), we next analysed RNA-Seq data to assess the expression levels of these genes. This revealed that only *STAT3* and *STAT5A*/*STAT5B* are expressed in mouse pBMDMs, while *STAT4* is not, ruling out this transcription factor as a mediator of macrophage reprogramming (Figure 3C, Table S1).

**Figure 3.**
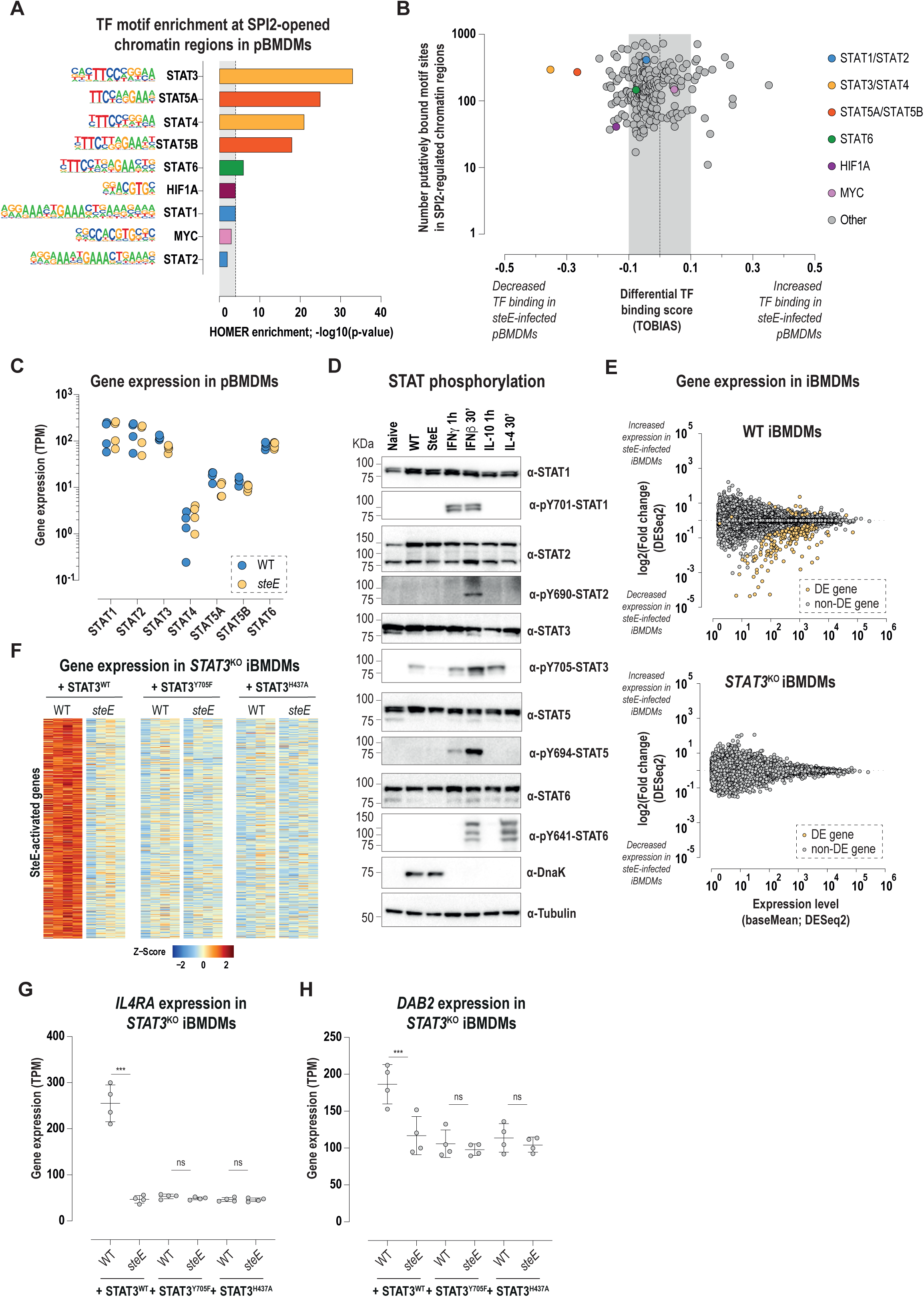
SteE-mediated transcriptional activation critically and specifically requires STAT3. **(A)** Enriched transcription factor motifs at SPI2-opened chromatin regions (see Figure 2A) were identified using HOMER software and the HOCOMOCO v11 MOUSE motif database. **(B)** Differential transcription factor footprinting analysis of combined SPI2-opened and SPI2-closed chromatin regions (see Figure 2A) in pBMDMs infected with *steE*-mutant ST19 *Salmonella* Typhimurium compared to those infected with the isogenic wild-type strain at 18 hours post-uptake. Footprinting was performed using TOBIAS (**<u>T</u>**ranscription factor **<u>O</u>**ccupancy prediction **<u>B</u>**<u>y</u> **I**nvestigation of **<u>A</u>**TAC-seq **<u>S</u>**ignal). The plot displays the differential transcription factor binding score (x-axis; calculated by TOBIAS) and the number of putative binding sites within SPI2-regulated chromatin regions for each transcription factor family (y-axis; calculated by TOBIAS). Selected transcription factor families are highlighted. **(C)** Gene expression levels of *STAT* family genes (RNA-Seq; see Figure 1) in pBMDMs infected with wild-type or *steE*-mutant ST19 *Salmonella* Typhimurium at 18 hours post-uptake. **(D)** Analysis of STAT protein phosphorylation in macrophages. pBMDMs were either untreated and unchallenged (naïve), infected with wild-type or *steE*-mutant ST19 *Salmonella* Typhimurium for 18 hours, or treated with 10 ng/mL IFN-γ, 20 ng/mL IL10 for 1 hour, or 1,000 U/mL IFN-β, 100 ng/mL IL4 for 30 minutes. Cell lysates were examined by immunoblotting for STAT protein expression and tyrosine phosphorylation, with DnaK and tubulin used as controls. Data shown are representative of three independent experiments. **(E)** Scatterplots illustrating differential gene expression between macrophages infected with wild-type ST19 *Salmonella* Typhimurium and an isogenic *steE*-mutant strain. Data are shown for wild type iBMDMs (upper panel) and *STAT3*^KO^ iBMDMs (bottom panel). The shrunken log_2_ fold change (y-axis, DESeq2), and the mean expression level (x-axis, DESeq2) are shown. Yellow dots highlight differentially expressed genes (adjusted p-value < 0.01, DESeq2). **(F)** Heatmap showing genes differentially activated (adjusted p-value < 0.01, DESeq2) in *STAT3*^KO^ iBMDMs complemented with STAT3^WT^, STAT3^Y705F^, or STAT3^H437A^ for macrophages infected with wild-type ST19 *Salmonella* Typhimurium or with an isogenic *steE*-mutant strain. Colours represent Z-scores, reflecting relative gene expression across all samples. **(G, H)** Gene expression levels for the anti-inflammatory marker genes *IL4RA* **(G)** and *DAB2* **(H)**, in *STAT3*^KO^ iBMDMs complemented with STAT3^WT^, STAT3^Y705F^, or STAT3^H43^ and infected with wild-type ST19 *Salmonella* Typhimurium or an isogenic *steE*-mutant strain. RNASeq was performed 18 hours post-uptake. Differential gene expression was analysed using DESeq2, with adjusted p-values shown (∗∗∗ adj. p < 0.001, ns = not significant).

We next measured STAT activation, which is regulated through tyrosine phosphorylation (Darnell et al., 1994; Ram et al., 1996; Shuai et al., 1993; Zhong et al., 1994), following an 18-hour infection of macrophages with *Salmonella* Typhimurium, a time point at which SPI2-driven transcriptional reprogramming is strongly evident (Stapels et al., 2018). Canonical stimuli were included in the analysis as positive controls. SteE translocation was specifically associated with phosphorylation of STAT3, with no phosphorylation of STAT2, STAT5 or STAT6 detected (Figure 3D). STAT1 phosphorylation was not detected in primary macrophages but was partially dependent on SteE in immortalised macrophages (Figure 3D and S3A).

To test the hypothesis that STAT3 is the primary transcription factor regulating macrophage polarisation induced by SteE, we generated *STAT3*^KO^ mouse immortalised bone marrow-derived macrophages (iBMDMs) (Figure S3B). As expected, in wild-type iBMDMs, *Salmonella* activated hundreds of genes in a SteE-dependent manner; however, in *STAT3*^KO^ macrophages, not a single gene was differentially expressed when comparing wild-type and *steE*-mutant infected cells (Figure 3E and S3C, Table S7). Consistent with a functional link between SteE-mediated STAT3 phosphorylation, its ability to activate an anti-inflammatory gene expression programme, and its role in stimulating chromatin remodelling in macrophages, SteE translocation failed to induce chromatin opening in *STAT3*^KO^ iBMDMs, in contrast to its effect in wild-type macrophages (Figure S3D, Table S8).

The ability of STAT3 to induce gene expression depends on: (1) activation through Y705 phosphorylation, which promotes dimerization and nuclear translocation, and (2) the capacity of phosphorylated STAT3 dimers to bind DNA at their specific motifs (Bromberg & Chen, 2001 and Darnell, 1997). Therefore, to further dissect the molecular mechanisms underlying SteE-mediated transcriptional activation, we introduced into *STAT3*^KO^ iBMDMs either a wild-type STAT3 (STAT3^WT^) or mutant versions of STAT3 (STAT3^Y705F^ and STAT3^H437A^). STAT3^Y705F^ cannot be activated via Y705 phosphorylation, and mutations of H437 result in significantly reduced STAT3 binding to DNA and are linked to Hyper-immunoglobulin E syndrome (Figure S3E-G and (Husby et al., 2012; Kaptein et al., 1996; Kasembeli et al., 2023; Minegishi et al., 2007; Mohr et al., 2013). In *STAT3*^KO^ iBMDMs expressing STAT3^WT^, pY705-STAT3 was detected in a SteE-dependent manner (Figure S3E-F), with SteE-mediated transcriptional activation fully restored, as demonstrated by the induction of canonical anti-inflammatory genes *IL4RA* and *DAB2* (Figure 3F-H, Table S9). In contrast, in macrophages expressing STAT3^Y705F^, which showed no pY705-STAT3 signal (Figure S3F), as expected (Kaptein et al., 1996; Minegishi et al., 2007; Mohr et al., 2013), or upon expression of STAT3^H437A^, which was phosphorylated (Figure S3F) but does not bind DNA (Kasembeli et al., 2023; Minegishi et al., 2007), SteE failed to induce SteE-activated genes including *IL4RA* or *DAB2* (Figs. 3F-H, Table S9). We therefore conclude that STAT3 is essential for SteE-induced gene activation.

Interestingly, STAT3 can heterodimerise with STAT1 (Zhong et al., 1994), and STAT1 phosphorylation was detected by others, and by our own studies in iBMDMs (Figure S4A). Therefore, to investigate the potential role of STAT1 further, we generated *STAT1*^KO^ iBMDMs (Figure S4A-B) and infected them with either wild-type or *steE*-mutant *Salmonella*. Surprisingly, but in line with our footprinting analysis (Figure 3B, Tables S5 and S6), the absence of STAT1 had no impact on SteE-mediated transcriptional activation (Figure S4C-E, Table S10). These findings suggest that, although SteE can induce STAT1 phosphorylation as previously reported (Stepien et al., 2024), STAT1 does not appear to play a biologically significant role in SteE-driven transcriptional reprogramming, at least in macrophages. Altogether, our results reveal that SteE-mediated transcriptional activation both critically and specifically requires STAT3.

### SteE translocation hyperactivates STAT3-target genes

Having identified STAT3 as the critical and specific mediator of SteE-induced macrophage transcriptional reprogramming, we next investigated whether SteE translocation simply phenocopies the effects of interleukin-10 (IL10) stimulation on macrophage phenotype (Hutchins et al., 2013; Moore et al., 2001). IL10 stimulation polarises macrophages towards an anti-inflammatory phenotype via phosphorylation and activation of STAT3 (Hutchins et al., 2013; Moore et al., 2001), and gene activation by IL10 stimulation is entirely dependent on this transcription factor (Figure S5A, Table S7).

To address this, we used RNA-Seq to compare the transcriptional activation triggered by SteE translocation – achieved by infecting macrophages with wild-type *Salmonella* Typhimurium versus an isogenic *steE*-mutant strain 18 hours after uptake – with the activation induced by stimulating naïve macrophages with a high dose of IL10 (20 ng/mL). Strikingly, even though both SteE translocation and IL10 treatment induced STAT3 phosphorylation (Figure S3B) and activated a similar set of genes associated with an anti-inflammatory macrophage phenotype, SteE translocation resulted in a significantly higher relative expression level of these genes compared to IL10 signalling (Figure 4A middle, 4B centre, Table S7). Furthermore, SteE translocation uniquely induced strong expression of numerous genes not activated by IL10 (Figure 4A top, 4B left, Table S7). Notably, the reverse is not observed: although IL10 activated hundreds of genes, these IL10-activated genes were expressed at much higher levels in *Salmonella*-infected macrophages at 18 hours post-uptake, regardless of infection with wild-type or *steE*-mutant bacteria (Figure 4A bottom, 4B right, Table S7). These findings demonstrate that SteE translocation drives transcriptional hyperactivation of STAT3-target genes, surpassing the effects of canonical IL10 signalling.

**Figure 4.**
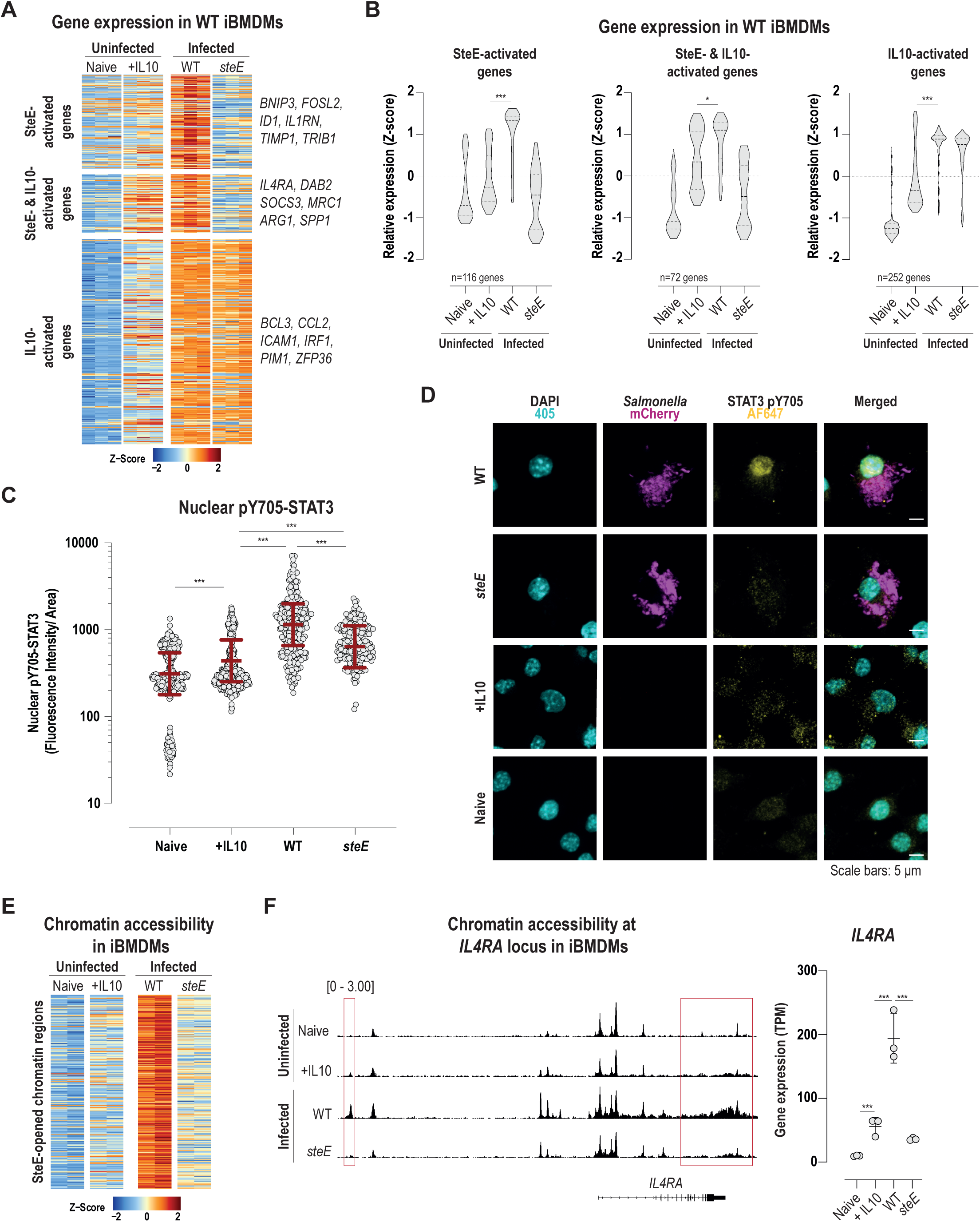
SteE hyperactivates STAT3-target genes. **(A-G)** Immortalized bone marrow-derived macrophages (iBMDMs) were infected with wild-type or *steE*-mutant ST19 *Salmonella* Typhimurium strains (12023/14028) or treated with 20 ng/mL IL10. Samples were analysed at 18 hours post-uptake or stimulation, unless otherwise specified. **(A)** Heatmap showing clusters of genes that are differentially activated (adjusted p-value < 0.01, DESeq2): only in iBMDMs infected with wild-type *Salmonella* Typhimurium compared to those infected with the *steE*-mutant strain (SteE-activated genes); in iBMDMs treated with IL10 compared to naïve (untreated and unchallenged) iBMDMs, and also in iBMDMs infected with wild-type *Salmonella* Typhimurium compared to those infected with the *steE*-mutant strain (SteE- and IL10-activated genes); and only in iBMDMs treated with IL10 compared to naïve (untreated and unchallenged) iBMDMs (IL10-activated genes). Colours represent Z-scores, indicating relative gene expression levels across all samples. **(B)** Violin plots visualising the relative expression levels (Z-scores) depicted in (A); FWER adjusted p-values calculated by GSEA software are shown. **(C)** Quantification of pY705-STAT3 in the cell nucleus following IL10 stimulation or wild-type of *steE*-mutant *Salmonella* infection (see (D)). Fluorescence intensity of pY705-STAT3 (AF647) in the cell nucleus (determined by DAPI stain) of naïve iBMDMs (untreated and unchallenged) and iBMDMs stimulated with IL10, or iBMDMs infected with wild type or *steE*-mutant ST19 *Salmonella* Typhimurium for 18 hours. Each point represents a single cell; red bars represent estimated marginal means with 95% confidence intervals for nuclear pY705-STAT3 fluorescence intensity across conditions based on a Gamma generalised linear mixed model with a log link, accounting for biological replicates (n=3 for each condition) as a random effect; *p*-values were calculated by Tukey-adjusted pairwise comparisons (∗∗∗ *p* < 0.001). **(D)** Nuclear localisation of pY705-STAT3. Representative confocal immunofluorescence microscopy images of iBMDMs stimulated as in **(C)**. Images represent nucleus (DAPI, Cyan), *Salmonella* (mCherry, Magenta) and pY705-STAT3 (AF647, yellow). Scale bars are 5 µm. Data shown is representative of three independent experiments. **(E)** Heatmap showing chromatin regions that are differentially opened (adjusted *p*-value < 0.1, limma) in iBMDMs infected with wild-type ST19 *Salmonella* Typhimurium compared to those infected with an isogenic *steE*-mutant. Colours represent Z-scores, reflecting relative chromatin accessibility levels across all samples. **(F)** Local chromatin accessibility profiles (left) and gene expression levels (right) for the anti-inflammatory marker gene *IL4RA* in naïve or IL10-treated iBMDMs and iBMDMs infected with wild type or *steE*-mutant ST19 *Salmonella* Typhimurium. Red boxes indicate regions encompassing selected differentially accessible chromatin regions (adjusted p-value < 0.1, limma); adjusted *p*-values for differential gene expression were calculated using DESeq2.

Given our finding that SteE induces transcriptional hyperactivation of STAT3-target genes, we next analysed STAT3 phosphorylation and nuclear localisation dynamics. Nuclear pY705-STAT3 levels in macrophages infected with wild-type *Salmonella* Typhimurium were significantly higher than those observed in either IL10-treated macrophages or macrophages infected with an isogenic *steE*-mutant strain (Figure 4C-D), mirroring the expression of STAT3-target genes observed under these conditions (Figure 4A-B). Furthermore, the dynamics of pY705-STAT3 signal in the macrophage upon IL10 treatment were consistent with negative feedback-loops, with Y705 phosphorylation levels oscillating over time (Figure S5B, Table S11). In contrast, infection with wild-type *Salmonella* Typhimurium induced progressive and sustained STAT3 phosphorylation (Figure S5B).

Finally, having established a functional link between SteE-mediated STAT3 phosphorylation, its activation of an anti-inflammatory gene expression programme and its role triggering chromatin remodelling in macrophages, we next investigated the extent of chromatin remodelling in both IL10 and *Salmonella*-infected conditions. Strikingly, IL10-mediated STAT3 phosphorylation did not induce robust opening of chromatin regions associated with SteE translocation (Figure 4E, Table S8). This is exemplified by the canonical anti-inflammatory gene *IL4RA*, where infection with wild-type *Salmonella* – but not IL10 stimulation – drives local alterations in chromatin accessibility with a concomitant hyperactivation of expression levels (Figure 4F, Tables S7 and S8).

Collectively, our findings demonstrate that SteE-induced STAT3 phosphorylation and DNA binding lead to hyperactivation of STAT3 target genes, surpassing the effects of canonical IL10 signalling. This hyperactivation correlates with elevated levels of phosphorylated STAT3 in the macrophage nucleus, which is associated with the opening of chromatin regions not opened upon physiological levels of STAT3 activation through endogenous cytokine signalling.

## DISCUSSION

In this study, we delineate the regulatory pathway linking the *Salmonella* Typhimurium effector SteE to host transcriptional reprogramming in macrophages, a key mechanism enabling bacterial survival, replication, and dissemination during bloodstream infection. Specifically, we show that SteE translocation induces M2-like macrophage responses by remodelling chromatin and opening cis-regulatory elements associated with anti-inflammatory gene activation. Although SteE has been proposed to exert transcriptionally repressive effects (Heyman et al., 2023), our results suggest these are secondary consequences of its broader capacity to reconfigure the macrophage gene expression landscape. We demonstrate that SteE-driven transcriptional reprogramming relies critically and specifically on STAT3 phosphorylation and DNA binding. Although both STAT3 and STAT1 are phosphorylated following SteE translocation, STAT1 phosphorylation is inconsistent, and its knockout does not affect SteE-induced gene activation in immortalised macrophages. This indicates that activation of multiple targets by SteE, such as STAT1 (Stepien et al., 2024), may reflect biological promiscuity, as observed with other effector proteins (Scott et al., 2017). Whereas previous studies have primarily raised concerns about effector promiscuity under artificial overexpression conditions compared to natural infection settings (Scott et al., 2017), our findings underscore the importance of genetic knockout studies for validating biologically relevant host pathways targeted by bacterial effectors, even in physiologically relevant infection models.

Most strikingly, we uncover that SteE induces hyperactivation of STAT3-target genes in macrophages, surpassing the effects of canonical IL10 signalling. This heightened activation is linked to increased levels of phosphorylated STAT3 in the macrophage nucleus and enhanced accessibility of cis-regulatory elements that are not opened by endogenous cytokine signalling. These findings underscore a key distinction between SteE-mediated STAT3 activation and IL10-driven signalling. IL10 signalling is initiated when IL10 binds to its receptor complex, comprising IL10R1 and IL10R2, associated with the Janus kinases JAK1 and TYK2, respectively (Finbloom’ & Winestock, 1995; Ho et al., 1995). Ligand binding promotes receptor dimerization, bringing the JAKs into proximity for cross-phosphorylation and activation (Lv et al., 2024). Activated JAKs phosphorylate tyrosine residues on the IL10R1 intracellular domain, generating docking sites for STAT3 via its SH2 domain (Weber-Nordtt et al., 1996). STAT3 is then phosphorylated at Y705, inducing dimerization through SH2-phosphotyrosine interactions. These STAT3 dimers translocate to the nucleus to regulate anti-inflammatory gene transcription (Lv et al., 2024). Critically, IL10 signalling is tightly regulated to prevent excessive activation: protein tyrosine phosphatases (e.g., SHP1) dephosphorylate JAKs and receptor phospho-tyrosines, disrupting STAT3 docking and activation; receptor complexes undergo clathrin-mediated endocytosis for degradation or recycling; and suppressors of cytokine signalling provide additional regulation with SOCS1 inhibiting JAK activity, while SOCS3 competes with STAT3 for receptor docking (Kim et al., 2018; Liau et al., 2018; Linossi & Nicholson, 2015). Together, these feedback loops ensure STAT3 activation is transient and proportional.

In contrast, SteE-mediated STAT3 activation bypasses many of the canonical negative feedback mechanisms by reprogramming GSK3 into a tyrosine kinase. Specifically, SteE interacts with GSK3, which induces the phosphorylation of SteE on several amino acids (Gibbs et al., 2020; Panagi et al., 2020). This step, which is critical for SteE activity (Panagi et al., 2020), results in GSK3-mediated phosphorylation of SteE at Y143 (Gibbs et al., 2020; Panagi et al., 2020), creating a pYxxQ motif that serves as a docking site for STAT3 (Gibbs et al., 2020). STAT3 is then phosphorylated on Y705 by GSK3, leading to dimerization, nuclear translocation and transcriptional activation (Panagi et al., 2020). Critically, unlike IL10 signalling, this pathway lacks a requirement for JAK kinases and therefore many of the known negative feedback mechanisms for STAT3 activation will be ineffective at dampening SteE-induced STAT3 phosphorylation (Gaggioli et al., 2024).

Although SteE plays a central role in STAT3 activation during *Salmonella* Typhimurium infection, infection itself also activates STAT3 independently of SteE. Lipopolysaccharide (LPS), a major component of the *Salmonella* outer membrane, triggers STAT3 activation in macrophages through TLR4-dependent signalling and subsequent IL6 and IL10 dependent feedback (Alonzi et al., 1998; Benkhart et al., 2000). SteE mediated activation of STAT3 via the reprogramming of GSK3 thus represents a highly specialised adaptation to modulate dynamic host signalling pathways. By bypassing regulatory constraints, SteE reprogrammes the macrophage gene regulatory network, enabling *Salmonella* Typhimurium to exploit a host anti-inflammatory transcription factor for its benefit. This mechanism drives the hyperactivation of STAT3 target genes, reconfigures the host cell phenotype, and enhances the ability of *Salmonella* to evade immune defences and establish systemic infection (Gibbs et al., 2020; Pham et al., 2020). Our study thus highlights the profound consequences of effector-mediated rewiring of host cell canonical signalling pathways, culminating in the enhanced pathogenesis of *Salmonella* (Gibbs et al., 2020; Pham et al., 2020). Even though we described the phenomenon in *Salmonella*, other bacterial effectors that activate host gene transcription via non-canonical pathways, such as *Bartonella* BepD, which hijacks the kinase c-Abl to activate STAT3 (Sorg et al., 2020), might similarly rewire host gene regulatory networks.

## STAR METHODS

### EXPERIMENTAL MODELS

#### Cell lines

Primary bone marrow derived macrophages (pBMDMs) were differentiated from monocytes isolated from the femur of 6 to 8-week old female C57BL/6 mice (Charles River). Red blood cells were lysed in 0.83% NH_4_Cl for 5 min. The remaining progenitor cells were cultured at 37°C and 5% CO_2_ in Dulbecco’s modified eagle medium with high glucose (DMEM; Sigma) containing 20% L929 (ATCC #CCL-1) conditioned media (LCM), 10% fetal bovine serum (FBS; Gibco), 10 mM HEPES (Sigma), 1 mM sodium pyruvate (Sigma), 0.05 mM beta-mercaptoethanol (Sigma), 100 U/ml penicillin (Sigma), and 100 µg/ml streptomycin (Sigma). Three days after isolation, medium was replaced. At day 7, differentiated pBMDMs were harvested and seeded.

Immortalized bone marrow derived macrophages (iBMDMs) were generated from female C57BL/6 mice using J2Cre virus to reduce the use of animals (Blasi et al., 1985). iBMDMs were cultured at 37 °C and 5% CO_2_ in DMEM; containing 20% LCM, 10% FBS, 10 mM HEPES, 1 mM sodium pyruvate and 0.05 mM beta-mercaptoethanol. pBMDMs and iBMDMs were seeded without LCM or antibiotics in tissue culture treated 6-well plates or 10 cm dishes. When indicated, 20 ng/ml recombinant murine IL10 (R & D) was added to the culture media for the designated duration.

HEK293ET cells (originating from a female fetus), were provided by Dr Felix Randow (MRC-LMB, Cambridge) and were maintained in DMEM with 10% FBS at 37 °C and 5% CO_2_.

#### Salmonella strains

*Salmonella enterica* serovar Typhimurium, gastrointestinal ST19 strain 12023/14028 and the clinical ST313 strain D23580, and its isogenic mutants, were used for all experiments. 12023/14028 *ssaV* mutant was from Prof. David Holden, Imperial College London and *steE* mutant was previously described (Panagi et al., 2020). D23580 was provided by Prof. Jay Hinton at the University of Liverpool. To generate the *ssaV* D23580 strain, homologous recombination was used as previously described (Datsenko & Wanner, 2000) and validated by genotyping PCR.

### METHODS DETAILS

#### iBMDM nucleofection

*STAT1*^KO^ and *STAT3*^KO^ iBMDMs were generated by nucleofection with px330-GFP plasmid (obtained from Dr Felix Randow). *STAT1* and *STAT3* gRNAs were designed using CHOPCHOP (Labun et al., 2019) and inserted in px330-GFP plasmids using BbsI (NEB) digestion followed by ligation with Taq DNA ligase (NEB) according to the manufacturer’s instructions. iBMDMs were nucleofected with 2.5 ng of *STAT1* or *STAT3* gRNA-containing px330-GFP plasmid using Amaxa Mouse Macrophage Nucleofection Kit (Lonza) following manufacturer’s instructions. Nucleofected iBMDMs were cultured as previously described. After 24 h, single GFP+ cells were sorted into 96 well plates using a BD FACS Aria and cultured under the same conditions for clonal expansion. Putative *STAT1*^KO^ or *STAT3*^KO^ clones were screened by immunoblot.

#### Retroviral particles production

M6p-puromycin plasmids (gift from Dr Felix Randow, MRC-LMB) were used to stably express wild type *STAT3* or *STAT3* mutants in iBMDMs. The *STAT3* open reading frame (UniProt: P40763) was amplified by PCR from HEK293ET cDNA and inserted in the m6p vector using NcoI and NotI. *STAT3* mutations were introduced by PCR-mediated mutagenesis. To generate retroviral particles, HEK293ET cells were transfected with m6p, pol/gag (Addgene plasmid 14887) and pMD2.VSVG (Addgene plasmid 12259) plasmids using Lipofectamine 2000 (Life Technologies) following manufacturer’s instructions. Media was replaced after 24 h, and 48 h later cell supernatants, containing retroviral particles, were collected and filtered through 0.45 µM filters before storage at -20 °C.

#### iBMDM transduction

Retroviral transduction was used to reconstitute *STAT3*^KO^ iBMDMs with *STAT3* WT or STAT3 mutants. Briefly, iBMDMs were seeded in 24 well plates and transduced with 1 mL of retroviral particle-containing cell supernatant and 8 μg/mL polybrene. Cells were centrifuged at 1000 x *g* at 30 °C for 1 h and incubated at 37 °C and 5% CO_2_ thereafter. After 72 h, transduced iBMDMs were selected using 3 µg/mL puromycin (Sigma).

#### Salmonella Infection

For fluorescence-activated cell sorting (FACS) strains containing the plasmid pFCcGi that constitutively expresses mCherry were used (Figueira et al., 2013). For all infections, bacteria were grown overnight in Luria broth (LB) medium at 37 °C and 200 rpm. Strains carrying pFCcGi were grown either with 50 µg/mL carbenicillin (STM12023/14028 background) or tetracycline (D23580 background) and mutant strains were grown with 50 µg/mL kanamycin. Bacteria were opsonised in 8% mouse serum (Sigma) for 20 min and then added to macrophages. Samples were centrifuged at 110 x *g* for 5 min, to synchronise bacteria uptake, and incubated for 30 min at 37 °C and 5% CO_2_. Non-phagocytosed bacteria were removed by washing twice with PBS, and macrophages were then cultured with 30 µg/mL gentamicin for 1 h and 10 µg/mL until further analysis at 37 °C and 5% CO_2_. Unless otherwise specified, macrophages were infected for 18 h.

#### Fluorescence-activated cell sorting (FACS)

Macrophages were washed in PBS twice and detached from the plates by scraping in cold PBS. Unfixed samples were sorted at 4 °C by a BD FACS Aria. Gating was used in all experiments to exclude apoptotic macrophages and doublets. Unless otherwise indicated in individual methods sections, for each sample, 10^5^ *Salmonella*-infected cells (mCherry+) and the same amount of naïve or IL10-treated macrophages were sorted for subsequent sequencing experiments.

#### RNA Sequencing

Uninfected macrophage samples were not challenged with *Salmonella*. For infected macrophage samples, *Salmonella* expressing the pFCcGi plasmid (i.e. arabinose inducible GFP, constitutive mCherry) (Figueira et al., 2013) were grown overnight in LB media without arabinose. Stationary phase bacteria were then opsonized with 8% mouse serum for 20 minutes and added to the BMDMs. For both primary and immortalised BMDMs (pBMDMs and iBMDMs) a MOI of 10 was used. At 30 min post-infection, macrophages were washed three times with PBS, and fresh BMDM medium containing gentamicin (20 µg/ml) was added to kill extracellular *Salmonella*. At 18 h post-infection, uninfected macrophage samples and *Salmonella*-infected gentamicin-treated macrophage samples were first washed with PBS three times and detached from the surface with cold PBS and scraping. Macrophages were then centrifuged at 4 °C at 300 x *g*, the supernatant discarded, and macrophages resuspended in cold sterile PBS. For infected macrophage samples, 50,000-100,000 macrophages containing *Salmonella* (i.e. mCherry^+^) were isolated by FACS. For uninfected macrophage samples, 50,000-100,000 macrophages were isolated by FACS without consideration of mCherry-status. For FACS isolation, apoptotic macrophages and doublets were excluded by gating, and the samples were sorted under continuous cooling to 4 °C by a BD Aria III into cold sterile PBS. Isolated macrophages in PBS were then centrifuged at 500 x *g* for 5 min at 4 °C, the supernatant was removed, and macrophage pellets were snap frozen in liquid nitrogen and stored at -80 °C until RNA was isolated. RNA was isolated using the Quick DNA-RNA Miniprep Kit following the manufacturer’s protocol. Following RNA isolation, mRNA-Seq libraries were generated using the NEBNext Ultra II Directional RNA Library Kit for Illumina (NEB E7765), the NEBNext Poly(A) mRNA Magnetic Isolation Module (NEB E7490), and the NEBNext Multiplex Oligos for Illumina (NEB E7335/E7500/E7710) following the manufacturer’s protocol. Libraries were sequenced on an Illumina NextSeq 2000 using paired-end sequencing.

For each biological replicate, raw fastq reads were trimmed using Trim Galore (version 0.6.4) with options --paired --illumina. Trimmed fastq files were then aligned to the mouse cDNA (mm10, UCSC) using the quant function of Kallisto (version 0.46.2) with option --rf-stranded.

For identification of genes with altered expression upon *Salmonella* infection and/or SPI2-mediated reprogramming in pBMDMs, gene-level abundance files generated by Kallisto (Bray et al., 2016) for each biological replicate were first loaded into R using tximport (version 1.24.0) and rhdf5 (version 2.40.0) to generate a count file (Soneson et al., 2016). The generated count file was then used to identify differentially expressed genes between pBMDMs infected with wild type *Salmonella* and pBMDMs infected with *ssaV Salmonella* using DESeq2 (Love et al., 2014) (version 1.36.0), with a model accounting for biological replicate batch effects (Replicate 1-4), *Salmonella* strain (12023/14028, D23580) and *Salmonella* genotype (wild type, *ssaV*). The generated count file was also used to identify differentially expressed genes between uninfected pBMDMs and pBMDMs infected with wild type *Salmonella* (12023 and D23580 background) using DESeq2 (version 1.36.0), with a model accounting for biological replicate batch effects (Replicate 1-4) and infection condition (uninfected, infected). In both instances, differentially expressed genes were defined as those with an adjusted p-value <0.01. The set of differentially expressed genes (between wild type *Salmonella*-infected pBMDMs and *ssaV Salmonella*-infected pBMDMs; and/or between uninfected pBMDMs and wild type *Salmonella*-infected pBMDMs) was used for subsequent principal component analysis and heatmap visualisation of DESeq2 (Love et al., 2014) (version 1.36.0) vst-normalised and limma (Ritchie et al., 2015) (version 3.52.4) batch-corrected count data. For Gene Set Enrichment Analysis (Subramanian et al., 2005)(GSEA, version 4.0.3), the stat value generated by DESeq2 following differential gene expression analysis (see above) was used to generate a pre-ranked gene list for testing the relative up-or down-regulation of either : 1) GSEA hallmark gene sets (Liberzon et al., 2015); or 2) GSEA hallmark gene sets (Liberzon et al., 2015), genes assigned to OCRs with statistically increased or decreased chromatin accessibility in pBMDMs infected with wild type *Salmonella* relative to pBMDMs infected with *ssaV Salmonella* (see ATAC-Seq methods below); all canonical pathway gene sets (Liberzon et al., 2011); and all gene ontology (GO) gene sets. For GSEA analysis, the weighted GSEA enrichment statistic was used to calculate the enrichment score and the FWER-adjusted p-value for all gene sets with between 15 and 2,000 genes.

For identification of genes with altered expression under various experimental conditions in wild type and *STAT3*^KO^, complemented *STAT3*^KO^, or *STAT1*^KO^ iBMDMs, gene-level abundance files generated by Kallisto for each biological replicate were first loaded into R using tximport (version 1.24.0) and rhdf5 (version 2.40.0) to generate a count file (Soneson et al., 2016). The generated count file was then used to identify differentially expressed genes using DESeq2 (version 1.36.0), with a model accounting for biological replicate batch effects and experimental condition. Differentially expressed genes were defined as those with an adjusted p-value <0.01. The set of differentially expressed genes was used for subsequent principal component analysis and heatmap visualisation of DESeq2 (version 1.36.0) vst-normalised and limma (version 3.52.4) batch-corrected count data.

Scripts to reproduce RNA-Seq associated analyses, figures and tables are provided in Supplementary File 1.

#### Assay for Transposase Accessible Chromatin (ATAC) Sequencing

primary and immortalised BMDMs (pBMDMs and iBMDMs), were infected as detailed in the RNA sequencing section. For infected macrophage samples, 50,000 macrophages containing *Salmonella* (i.e. mCherry^+^) were isolated by FACS as outlined above. For uninfected macrophage samples, 50,000 macrophages were isolated by FACS without consideration of mCherry-status. Following isolation of the macrophages, ATAC-Seq libraries were immediately generated using the ATAC-Seq Kit from Active Motif (Catalogue no. 53150), following the manufacturer’s protocol. Libraries were sequenced on an Illumina NextSeq 2000 using paired-end sequencing.

For each biological replicate, raw fastq reads were trimmed using Trim Galore (version 0.6.4) with options --paired --nextera. Trimmed fastq files were then aligned to the mouse genome (mm10, UCSC) using bowtie2 (version 2.4.1) with options --very-sensitive -X 1000 - k 1. The aligned bam file was then sorted using the SortSam function of Picard (version 2.22.3) with options SORT_ORDER=coordinate VALIDATION_STRINGENCY=LENIENT, and then deduplicated using the MarkDuplicates function of Picard (version 2.22.3) with options VALIDATION_STRINGENCY=LENIENT REMOVE_DUPLICATES=true. Following this, reads overlapping chrM and the mm10 blacklisted regions (Amemiya et al., 2019) were removed using the intersectBed function of bedtools (Quinlan & Hall, 2010) (version 2.30.0) with option -v, and the subsequent deduplicated and cleaned bam file was sorted and indexed using the sort and index functions of samtools (Li et al., 2009) (version 1.9). The sorted, cleaned and deduplicated bam files for each biological replicate were then used to call open chromatin regions (OCRs) using the callpeak function of MACS2 (Feng et al., 2012) (version 2.2.7.1) with options -f BAMPE -g mm --keep-dup all -p 0.01 --nolambda -- bdg --SPMR. To identify reproducible OCRs, the sorted, cleaned and deduplicated bam files corresponding to individual biological replicates were pooled using the merge function of samtools (version 1.9), and this pooled bam file was then sorted and indexed using the sort and index functions of samtools (version 1.9). The pooled bam file was then used to call “pooled OCRs” using the callpeak function of MACS2 (version 2.2.7.1) with options -f BAMPE -g mm --keep-dup all -p 0.01 --nolambda --bdg –SPMR. Finally, the intersectBed function of bedtools (v2.30.0) was used to select reproducible OCRs, defined here as “pooled OCRs” that overlap OCRs also identified in every biological replicate; where only two biological replicates were available (i.e. for iBMDM samples), the option -F 0.50 was used to provide increased stringency with regards to reproducible OCRs.

For identification of OCRs with altered accessibility upon *Salmonella* infection and/or SPI2-mediated reprogramming in pBMDMs, a consensus open chromatin region (OCR) BED file was first generated by concatenating reproducible OCRs from uninfected pBMDMs, pBMDMs infected with wild type *Salmonella* (12023/14028 and D23580 background) and pBMDMs infected with *ssaV Salmonella* (12023/14028 and D23580 background), and then merging overlapping OCRs with the merge function of bedtools (version 2.30.0), with no additional options. This consensus OCR BED file was then converted to SAF format using custom bash script. This SAF file was used to count reads aligning to consensus OCRs from uninfected pBMDMs, pBMDMs infected with wild type *Salmonella* (12023/14028 and D23580 background), pBMDMs infected with *ssaV Salmonella* (12023/14028 and D23580 background) and pBMDMs infected with *steE Salmonella* (STM12023/14028 background) using the featureCounts function of Subread (Liao, et al., 2014) (version 2.0.1) with options - F SAF -O --fracOverlap 0.2 -p. The generated count file was used to identify differentially accessible OCRs between pBMDMs infected with wild type *Salmonella* and pBMDMs infected with *ssaV Salmonella* using DESeq2 (Love, et al., 2014) (version 1.36.0), with a model accounting for biological replicate batch effects (Replicate 1-4), *Salmonella* strain (12023/14028, D23580) and *Salmonella* genotype (wild type, *ssaV*). The generated count file was also used to identify differentially accessible OCRs between uninfected pBMDMs and pBMDMs infected with wild type *Salmonella* (12023/14028 and D23580 background) using DESeq2 (version 1.36.0), with a model accounting for biological replicate batch effects (Replicate 1-4) and infection condition (uninfected, infected). In both instances, differentially accessible OCRs were defined as those with an adjusted p-value <0.1. The set of differentially accessible OCRs (between wild type *Salmonella*-infected pBMDMs and *ssaV Salmonella*-infected pBMDMs; and/or between uninfected pBMDMs and wild type Salmonella-infected pBMDMs) was used for subsequent principal component analysis and heatmap visualisation of vst-normalised and limma (Ritchie, et al., 2015) (version 3.52.4) batch-corrected count data. To identify enriched transcription factor motifs at SPI2-opened chromatin, the findMotifsGenome.pl function of HOMER (version 4.11) was used (Duttke, et al., 2019), with OCRs specifically showing increased accessibility (based on DESeq2 analysis, see above) between pBMDMs infected with wild type *Salmonella* and pBMDMs infected with *ssaV Salmonella* as input, all consensus OCRs as background, the HOCOMOCO v11 MOUSE motifs as known motif database, and options -mask -size given. GREAT (McLean, et al., 2010; Tanigawa, et al., 2022)(version 4.0.4) was used to assign OCRs with statistically increased or decreased chromatin accessibility in pBMDMs infected with wild type *Salmonella* relative to pBMDMs infected with *ssaV Salmonella* to nearby genes using the basal plus extension association model with options Proximal: TSS -5 kb/+1 kb, Distal: TSS +/-1,000 kb and mm10 curated regulatory domains included.

To identify differential transcription factor activity at OCRs showing differential accessibility in pBMDMs infected with wild type *Salmonella* and pBMDMs infected with *ssaV Salmonella*, transcription factor footprinting analysis was carried out using TOBIAS (Bentsen et al., 2020) (version 0.12.12). In brief, the ATACorrect function of TOBIAS was first used to correct for the Tn5 transposase insertion bias across consensus OCRs for each pooled bam file associated with a specific experimental condition (i.e. wild type, *ssaV* or *steE Salmonella*-infected pBMDMs). The corresponding bias-corrected single base pair cutsite Bigwig tracks were then used to calculate a continuous transcription factor footprinting score for each condition across all consensus OCRs using the FootprintScores function of TOBIAS. Finally, these transcription factor footprinting scores were used to predict differential transcription factor binding at OCRs specifically showing differential accessibility (based on DESeq2 analysis, see above) between pBMDMs infected with wild type *Salmonella* and pBMDMs infected with *ssaV Salmonella* using the BINDetect function of TOBIAS with option --motifs H11_MOUSE_mono_jaspar_format.txt (i.e. HOCOMOCO v11 MOUSE motifs in JASPER format).

To identify OCRs with altered accessibility because of SteE-mediated reprogramming in wild type iBMDMs, a consensus open chromatin region (OCR) BED file was first generated by concatenating and merging reproducible OCRs from wild type iBMDMs infected with wild type or *steE Salmonella* (12023/14028 strain). This consensus OCR BED file was then converted to SAF format using custom bash script. This SAF file was used to count reads aligning to consensus OCRs from uninfected naïve iBMDMs (wild type and *STAT3*^KO^ genotypes), uninfected IL10-treated iBMDMs (wild type and *STAT3*^KO^ genotypes), wild type and *STAT3*^KO^ iBMDMs infected with wild type *Salmonella*, and wild type and *STAT3*^KO^ iBMDMs infected with *steE Salmonella*. Read counts were calculated using the featureCounts function of Subread (Liao et al., 2014)(version 2.0.1) with options -F SAF -O - -fracOverlap 0.2 -p. The generated count file was then used to identify the top 20,000 variable OCRs, which were subsequently used to carry out differential accessibility analysis between wild type and *steE Salmonella*-infected wild type iBMDMs using limma (version 3.52.4), with a model accounting for biological replicate batch effects and experimental condition. Of note, limma software was used in this instance instead of DESeq2, as benchmarking experiments have previously shown that limma has markedly increased sensitivity to identify differentially accessible OCRs when only two biological replicates per condition are available and regions of interest are known to have relatively low ATAC-Seq signal (Gontarz et al., 2020), conditions that were relevant for our analysis. An adjusted p-value cutoff of < 0.1 was then used to identify differentially accessible OCRs for visualisation of DESeq2 (version 1.36.0) vst-normalised and limma (version 3.52.4) batch-corrected count data.

Scripts to reproduce ATAC-Seq associated analyses, figures and tables are provided in Supplementary File 1.

#### Microscopy

Macrophages were grown on glass coverslips, infected as described above, washed twice with PBS and fixed in 4% paraformaldehyde in PBS for 20 min. Cells were then permeabilized and blocked in 0.1% Triton X-100 and 10% horse serum in PBS (blocking solution) for 30 min and stained with anti-pY705 STAT3 antibody in blocking solution for 1 h at room temperature. After three washes with PBS, cells were incubated with AlexaFluor 647 anti-rabbit secondary antibody (Alexa-Flour, Invitrogen) and 0.5 mg/mL diamidino-2-phenylindole (DAPI, Invitrogen) in blocking solution for 1 h at room temperature. Samples were then washed three times with PBS and once with dH_2_0, mounted on glass slides with Aqua-Poly/Mount (Polysciences, Inc.) and imaged on an LSM 710 inverted confocal microscope (Zeiss) using a x 63, 1.4 numerical aperture objective. Confocal images were analysed using ImageLab software. The “ROI Manager” tool was used to define nuclear regions of interest (ROIs) based on DAPI stain. ImageJ “Analyze Particles” function was applied to measure the nuclear ROIs areas and to quantify the mean fluorescence intensity of AF647, labelling pY705-STAT3, within these nuclear ROIs. Estimated marginal means with 95% confidence intervals for nuclear pY705-STAT3 fluorescence intensity across conditions were calculated using a Gamma generalised linear mixed model with a log link, accounting for biological replicates (n=3 for each condition) as a random effect, and *p*-values were calculated by Tukey-adjusted pairwise comparisons.

#### SDS-PAGE and Immunoblotting

To prepare whole cell lysates, macrophages were washed once in PBS and lysed in Lumier’s Lysis buffer (20 mM Tris HCl pH7.4, 150 mM NaCl, 10% glycerol, 0.3% Triton X- 100) supplemented with proteases and phosphatases inhibitors (cOmplete EDTA-free Protease Inhibitor Cocktail and PhosSTOP, Roche) for 30 min on ice and sonicated.

Whole lysates were denatured at 95 °C for 5 min in SDS loading buffer. Lysates were separated by SDS-PAGE using 10 % polyacrylamide denaturing gels, transferred to 0.2 µm PVDF membranes (Millipore) and visualized by immunoblotting using ECL detection reagents (Dako) on an iBright 1500 Imager (Thermofisher).

### STATISTICAL ANALYSIS

Statistical analyses associated with RNA-Seq, ATAC-Seq and microscopy datasets are detailed in their associated methods sections and/or figure legends and relevant scripts are provided in Supplementary File 1. For other statistical analyses, GraphPad Prism software was used to test statistical significance of the data. The statistical test performed and the number of replicates for each experiment are specified in the figure legends.

## Data Availability Section

### Lead contact

Further information and requests for resources and reagents should be directed to and will be fulfilled by the Lead Contact, Teresa L. M. Thurston.

### Materials availability

Plasmids and bacterial strains generated in this study are available upon request to the Lead Contact.

### Data and code availability

All data reported in this paper will be shared by the lead contact upon request.

## Acknowledgements and Funding

We thank members of the Thurston and Hill laboratories for scientific discussion and reading of the manuscript; Jay Hinton for providing the clinical isolate used in this study; Felix Randow for plasmids and cell lines; David Holden for *Salmonella* strains, Jessica Rowley and Larissa Zarate for Fluorescence Activated Cell Sorting and Katerina Rekopoulou at the Imperial BRC Genomics Facility for running the next-generation sequencing.

Research reported in this publication was funded by: a Biotechnology and Biological Sciences Research Council (BBSRC) David Phillips Fellowship (BB/R011834/1) awarded to TLMT, which also supported IP; a Medical Research Council (MRC) grant MR/V031058/1 awarded to TLMT, which funded IDDO; an Engineering and Physical Sciences Research Council (EPSRC) grant EP/X02377X/1, underwriting European Research Council Starting Grant, Re-kin awarded to TLMT, which supported IP and POS; an MRC DTP Studentship (MR/N014103/1) awarded to IP, an Imperial College Research Fellowship and Wellcome Trust Career Development Award (226546/Z/22/Z) to PWSH; an EMBO Scientific Exchange Grant 9578 and the NAWA International Interdisciplinary Doctoral School at the HEART of BioBased BPI/STE2021/1/00008/U/00001 to AS; The Japan Society for the Promotion of Science (JSPS) KAKENHI grants JP 21J12222 to SS.

For open access, the author has applied a CC BY public copyright license to any author-accepted manuscript version arising from this submission.

## Author contributions

Conceptualization: PWSH and TLMT; Methodology: IDDO, POS, PWSH and TLMT; Investigation: IDDO, SS, AS, IP, POS and PWSH; Writing – Original Draft: IDDO, TLMT and PWSH; Writing – Review & Editing: IDDO, PWSH and TLMT; Funding Acquisition: TLMT and PWSH; Supervision: MO, KG, PWSH and TLMT.

## Disclosure and Competing Interests

The authors declare no competing interests.

## Notes

### Competing Interest Statement

The authors have declared no competing interest.

## REFERENCES

1. Alonzi, T., Fattori, E., Cappelletti, M., Ciliberto, G., & Poli, V. (1998). Impaired STAT3 activation following localized inflammatory stimulus in IL-6-deficient mice. Cytokine, 10(1), 13–18. 10.1006/CYTO.1997.0250

2. Amemiya, H. M., Kundaje, A., & Boyle, A. P. (2019). The ENCODE Blacklist: Identification of Problematic Regions of the Genome. Scientific Reports 2019 9:1, 9(1), 1–5. 10.1038/s41598-019-45839-z

3. Benkhart, E. M., Siedlar, M., Wedel, A., Werner, T., & Ziegler-Heitbrock, H. W. L. (2000). Role of Stat3 in Lipopolysaccharide-Induced IL-10 Gene Expression. The Journal of Immunology, 165(3), 1612–1617. 10.4049/JIMMUNOL.165.3.1612

4. Bentsen, M., Goymann, P., Schultheis, H., Klee, K., Petrova, A., Wiegandt, R., Fust, A., Preussner, J., Kuenne, C., Braun, T., Kim, J., & Looso, M. (2020). ATAC-seq footprinting unravels kinetics of transcription factor binding during zygotic genome activation. Nature Communications 2020 11:1, 11(1), 1–11. 10.1038/s41467-020-18035-1

5. Blasi, E., Mathieson, B. J., Varesio, L., Cleveland, J. L., Borchert, P. A., & Rapp, U. R. (1985). Selective immortalization of murine macrophages from fresh bone marrow by a raf/myc recombinant murine retrovirus. Nature 1985 318:6047, 318(6047), 667–670. 10.1038/318667a0

6. Bray, N. L., Pimentel, H., Melsted, P., & Pachter, L. (2016). Near-optimal probabilistic RNA-seq quantification. Nature Biotechnology, 34(5), 525–527. 10.1038/NBT.3519

7. Buenrostro, J. D., Giresi, P. G., Zaba, L. C., Chang, H. Y., & Greenleaf, W. J. (2013). Transposition of native chromatin for fast and sensitive epigenomic profiling of open chromatin, DNA-binding proteins and nucleosome position. Nature Methods 2013 10:12, 10(12), 1213–1218. 10.1038/nmeth.2688

8. Darnell, J. E., Kerr, lan M., & Stark, G. R. (1994). Jak-STAT Pathways and Transcriptional Activation in Response to IFNs and Other Extracellular Signaling Proteins. Science, 264(5164), 1415–1421. 10.1126/SCIENCE.8197455

9. Datsenko, K. A., & Wanner, B. L. (2000). One-step inactivation of chromosomal genes in Escherichia coli K-12 using PCR products. Proceedings of the National Academy of Sciences of the United States of America, 97(12), 6640–6645. 10.1073/PNAS.120163297/ASSET/103C1B8D-D302-4337-8E67-9F9084156407/ASSETS/GRAPHIC/PQ1201632006.JPEG

10. Duttke, S. H., Chang, M. W., Heinz, S., & Benner, C. (2019). Identification and dynamic quantification of regulatory elements using total RNA. Genome Research, 29(11), 1836–1846. 10.1101/GR.253492.119

11. Feng, J., Liu, T., Qin, B., Zhang, Y., & Liu, X. S. (2012). Identifying ChIP-seq enrichment using MACS. Nature Protocols 2012 7:9, 7(9), 1728–1740. 10.1038/nprot.2012.101

12. Fields, P. I., Swanson, R. V., Haidaris, C. G., & Heffron, F. (1986). Mutants of Salmonella typhimurium that cannot survive within the macrophage are avirulent. Proceedings of the National Academy of Sciences, 83(14), 5189–5193. 10.1073/PNAS.83.14.5189

13. Figueira, R., Watson, K. G., Holden, D. W., & Helaine, S. (2013). Identification of Salmonella Pathogenicity Island-2 Type III Secretion System Effectors Involved in Intramacrophage Replication of S. enterica Serovar Typhimurium: Implications for Rational Vaccine Design. MBio, 4(2), e00065–13. 10.1128/MBIO.00065-13

14. Gaggioli, M. R., Jones, A. G., Panagi, I., Washington, E. J., Loney, R. E., Muench, J. H., Brennan, R. G., Thurston, T. L. M., & Ko, D. C. (2024). A single amino acid in the Salmonella effector SarA/SteE triggers supraphysiological activation of STAT3 for anti-inflammatory target gene expression. BioRxiv, 2024.02.14.580367. 10.1101/2024.02.14.580367

15. Gibbs, K. D., Washington, E. J., Jaslow, S. L., Bourgeois, J. S., Foster, M. W., Guo, R., Brennan, R. G., & Ko, D. C. (2020). The Salmonella Secreted Effector SarA/SteE Mimics Cytokine Receptor Signaling to Activate STAT3. Cell Host & Microbe, 27(1), 129. 10.1016/J.CHOM.2019.11.012

16. Gontarz, P., Fu, S., Xing, X., Liu, S., Miao, B., Bazylianska, V., Sharma, A., Madden, P., Cates, K., Yoo, A., Moszczynska, A., Wang, T., & Zhang, B. (2020). Comparison of differential accessibility analysis strategies for ATAC-seq data. Scientific Reports 2020 10:1, 10(1), 1–13. 10.1038/s41598-020-66998-4

17. Heyman, O., Yehezkel, D., Mattioli, C. C., Blumberger, N., Rosenberg, G., Solomon, A., Hoffman, D., Ben-Moshe, N. B., & Avraham, R. (2023). Paired single-cell host profiling with multiplex-tagged bacterial mutants reveals intracellular virulence-immune networks. Proceedings of the National Academy of Sciences of the United States of America, 120(28), e2218812120. 10.1073/PNAS.2218812120/SUPPL_FILE/PNAS.2218812120.SAPP.PD F

18. Husby, J., Todd, A. K., Haider, S. M., Zinzalla, G., Thurston, D. E., & Neidle, S. (2012). Molecular dynamics studies of the STAT3 homodimer:DNA complex: Relationships between STAT3 mutations and protein-DNA recognition. Journal of Chemical Information and Modeling, 52(5), 1179–1192. 10.1021/CI200625Q/SUPPL_FILE/CI200625Q_SI_001.PDF

19. Hutchins, A. P., Diez, D., & Miranda-Saavedra, D. (2013). The IL-10/STAT3-mediated anti-inflammatory response: recent developments and future challenges. Briefings in Functional Genomics, 12(6), 489. 10.1093/BFGP/ELT028

20. Jaslow, S. L., Gibbs, K. D., Fricke, W. F., Wang, L., Pittman, K. J., Mammel, M. K., Thaden, J. T., Fowler, V. G., Hammer, G. E., Elfenbein, J. R., & Ko, D. C. (2018). Salmonella Activation of STAT3 Signaling by SarA Effector Promotes Intracellular Replication and Production of IL-10. Cell Reports, 23(12), 3525–3536. 10.1016/J.CELREP.2018.05.072

21. Jennings, E., Thurston, T. L. M., & Holden, D. W. (2017). Salmonella SPI-2 Type III Secretion System Effectors: Molecular Mechanisms And Physiological Consequences. Cell Host & Microbe, 22(2), 217–231. 10.1016/J.CHOM.2017.07.009

22. Kaptein, A., Paillard, V., & Saunders, M. (1996). Dominant negative Stat3 mutant inhibits interleukin-6-induced Jak-STAT signal transduction. Journal of Biological Chemistry, 271(11), 5961–5964. 10.1074/jbc.271.11.5961

23. Kasembeli, M. M., Kaparos, E., Bharadwaj, U., Allaw, A., Khouri, A., Acot, B., & Tweardy, D. J. (2023). Aberrant function of pathogenic STAT3 mutant proteins is linked to altered stability of monomers and homodimers. Blood, 141(12), 1411–1424. 10.1182/BLOOD.2021015330

24. Kulakovskiy, I. V., Vorontsov, I. E., Yevshin, I. S., Sharipov, R. N., Fedorova, A. D., Rumynskiy, E. I., Medvedeva, Y. A., Magana-Mora, A., Bajic, V. B., Papatsenko, D. A., Kolpakov, F. A., & Makeev, V. J. (2018). HOCOMOCO: towards a complete collection of transcription factor binding models for human and mouse via large-scale ChIP-Seq analysis. Nucleic Acids Research, 46(D1), D252–D259. 10.1093/NAR/GKX1106

25. Labun, K., Montague, T. G., Krause, M., Torres Cleuren, Y. N., Tjeldnes, H., & Valen, E. (2019). CHOPCHOP v3: expanding the CRISPR web toolbox beyond genome editing. Nucleic Acids Research, 47(W1), W171–W174. 10.1093/NAR/GKZ365

26. Lawrence, T., & Natoli, G. (2011). Transcriptional regulation of macrophage polarization: enabling diversity with identity. Nature Reviews Immunology 2011 11:11, 11(11), 750–761. 10.1038/nri3088

27. Li, H., Handsaker, B., Wysoker, A., Fennell, T., Ruan, J., Homer, N., Marth, G., Abecasis, G., & Durbin, R. (2009). The Sequence Alignment/Map format and SAMtools. *Bioinformatics (Oxford*, England), 25(16), 2078–2079. 10.1093/BIOINFORMATICS/BTP352

28. Liao, Y., Smyth, G. K., & Shi, W. (2014). featureCounts: an efficient general purpose program for assigning sequence reads to genomic features. *Bioinformatics (Oxford*, England), 30(7), 923–930. 10.1093/BIOINFORMATICS/BTT656

29. Liberzon, A., Birger, C., Thorvaldsdóttir, H., Ghandi, M., Mesirov, J. P., & Tamayo, P. (2015). The Molecular Signatures Database (MSigDB) hallmark gene set collection. Cell Systems, 1(6), 417–425. 10.1016/J.CELS.2015.12.004

30. Liberzon, A., Subramanian, A., Pinchback, R., Thorvaldsdóttir, H., Tamayo, P., & Mesirov, J. P. (2011). Molecular signatures database (MSigDB) 3.0. Bioinformatics (Oxford, England), 27(12), 1739–1740. 10.1093/BIOINFORMATICS/BTR260

31. Love, M. I., Huber, W., & Anders, S. (2014). Moderated estimation of fold change and dispersion for RNA-seq data with DESeq2. Genome Biology, 15(12). 10.1186/S13059-014-0550-8

32. Lv, Y., Qi, J., Babon, J. J., Cao, L., Fan, G., Lang, J., Zhang, J., Mi, P., Kobe, B., & Wang, F. (2024). The JAK-STAT pathway: from structural biology to cytokine engineering. Signal Transduction and Targeted Therapy 2024 9:1, 9(1), 1–37. 10.1038/s41392-024-01934-w

33. McLean, C. Y., Bristor, D., Hiller, M., Clarke, S. L., Schaar, B. T., Lowe, C. B., Wenger, A. M., & Bejerano, G. (2010). GREAT improves functional interpretation of cis-regulatory regions. Nature Biotechnology, 28(5), 495–501. 10.1038/NBT.1630

34. Minegishi, Y., Saito, M., Tsuchiya, S., Tsuge, I., Takada, H., Hara, T., Kawamura, N., Ariga, T., Pasic, S., Stojkovic, O., Metin, A., & Karasuyama, H. (2007). Dominant-negative mutations in the DNA-binding domain of STAT3 cause hyper-IgE syndrome. Nature 2007 448:7157, 448(7157), 1058–1062. 10.1038/nature06096

35. Mohr, A., Fahrenkamp, D., Rinis, N., & Müller-Newen, G. (2013). Dominant-negative activity of the STAT3-Y705F mutant depends on the N-terminal domain. Cell Communication and Signaling, 11(1), 1–12. 10.1186/1478-811X-11-83/FIGURES/6

36. Monack, D. M. (2013). Helicobacter and Salmonella Persistent Infection Strategies. Cold Spring Harbor Perspectives in Medicine, 3(12), a010348. 10.1101/CSHPERSPECT.A010348

37. Moore, K. W., De Waal Malefyt, R., Coffman, R. L., & O’Garra, A. (2001). Interleukin-10 and the interleukin-10 receptor. Annual Review of Immunology, 19(Volume 19, 2001), 683–765. 10.1146/ANNUREV.IMMUNOL.19.1.683/CITE/REFWORKS

38. Panagi, I., Jennings, E., Zeng, J., Günster, R. A., Stones, C. D., Mak, H., Jin, E., Stapels, D. A. C., Subari, N. Z., Pham, T. H. M., Brewer, S. M., Ong, S. Y. Q., Monack, D. M., Helaine, S., & Thurston, T. L. M. (2020). Salmonella Effector SteE Converts the Mammalian Serine/Threonine Kinase GSK3 into a Tyrosine Kinase to Direct Macrophage Polarization. Cell Host & Microbe, 27(1), 41–53.e6. 10.1016/J.CHOM.2019.11.002

39. Pham, T. H. M., Brewer, S. M., Thurston, T., Massis, L. M., Honeycutt, J., Lugo, K., Jacobson, A. R., Vilches-Moure, J. G., Hamblin, M., Helaine, S., & Monack, D. M. (2020). Salmonella-Driven Polarization of Granuloma Macrophages Antagonizes TNF-Mediated Pathogen Restriction during Persistent Infection. Cell Host & Microbe, 27(1), 54–67.e5. 10.1016/J.CHOM.2019.11.011

40. Pham, T. H. M., Xue, Y., Brewer, S. M., Bernstein, K. E., Quake, S. R., & Monack, D. M. (2023). Single-cell profiling identifies ACE+ granuloma macrophages as a nonpermissive niche for intracellular bacteria during persistent Salmonella infection. Science Advances, 9(1). 10.1126/SCIADV.ADD4333/SUPPL_FILE/SCIADV.ADD4333_SM.PDF

41. Quinlan, A. R., & Hall, I. M. (2010). BEDTools: a flexible suite of utilities for comparing genomic features. *Bioinformatics (Oxford*, England), 26(6), 841–842. 10.1093/BIOINFORMATICS/BTQ033

42. Ram, P. A., Park, S. H., Choi, H. K., & Waxman, D. J. (1996). Growth hormone activation of Stat 1, Stat 3, and Stat 5 in rat liver: Differential kinetics of hormone desensitization and growth hormone stimulation of both tyrosine phosphorylation and serine/threonine phosphorylation. Journal of Biological Chemistry, 271(10), 5929–5940. 10.1074/JBC.271.10.5929/ASSET/77276BAF-7750-456A-B60F-D6F04256AF15/MAIN.ASSETS/GR5.JPG

43. Richter-Dahlfors, A., Buchan, A. M. J., & Finlay, B. B. (1997). Murine Salmonellosis Studied by Confocal Microscopy: Salmonella typhimurium Resides Intracellularly Inside Macrophages and Exerts a Cytotoxic Effect on Phagocytes In Vivo. Journal of Experimental Medicine, 186(4), 569–580. 10.1084/JEM.186.4.569

44. Ritchie, M. E., Phipson, B., Wu, D., Hu, Y., Law, C. W., Shi, W., & Smyth, G. K. (2015). limma powers differential expression analyses for RNA-sequencing and microarray studies. Nucleic Acids Research, 43(7), e47. 10.1093/NAR/GKV007

45. Salcedo, S. P., Noursadeghi, M., Cohen, J., & Holden, D. W. (2001). Intracellular replication of Salmonella typhimurium strains in specific subsets of splenic macrophages in vivo. Cellular Microbiology, 3(9), 587–597. 10.1046/J.1462-5822.2001.00137.X

46. Scott, N. E., Giogha, C., Pollock, G. L., Kennedy, C. L., Webb, A. I., Williamson, N. A., Pearson, J. S., & Hartland, E. L. (2017). The bacterial arginine glycosyltransferase effector NleB preferentially modifies Fas-associated death domain protein (FADD). The Journal of Biological Chemistry, 292(42), 17337. 10.1074/JBC.M117.805036

47. Shuai, K., Stark, G. R., Kerr, I. M., & Darnell, J. E. (1993). A Single Phosphotyrosine Residue of Stat91 Required for Gene Activation by Interferon-γ. Science, 261(5129), 1744–1746. 10.1126/SCIENCE.7690989

48. Soneson, C., Love, M. I., & Robinson, M. D. (2016). Differential analyses for RNA-seq: transcript-level estimates improve gene-level inferences. F1000Research 2015 4:1521, 4, 1521. 10.12688/f1000research.7563.1

49. Sorg, I., Schmutz, C., Lu, Y. Y., Fromm, K., Siewert, L. K., Bögli, A., Strack, K., Harms, A., & Dehio, C. (2020). A Bartonella Effector Acts as Signaling Hub for Intrinsic STAT3 Activation to Trigger Anti-inflammatory Responses. Cell Host & Microbe, 27(3), 476–485.e7. 10.1016/J.CHOM.2020.01.015

50. Stanaway, J. D., Parisi, A., Sarkar, K., Blacker, B. F., Reiner, R. C., Hay, S. I., Nixon, M. R., Dolecek, C., James, S. L., Mokdad, A. H., Abebe, G., Ahmadian, E., Alahdab, F., Alemnew, B. T., Alipour, V., Bakeshei, F. A., Animut, M. D., Ansari, F., Arabloo, J., … Crump, J. A. (2019). The global burden of non-typhoidal salmonella invasive disease: a systematic analysis for the Global Burden of Disease Study 2017. The Lancet Infectious Diseases, 19(12), 1312–1324. 10.1016/S1473-3099(19)30418-9

51. Stapels, D. A. C., Hill, P. W. S., Westermann, A. J., Fisher, R. A., Thurston, T. L., Saliba, A. E., Blommestein, I., Vogel, J., & Helaine, S. (2018). Salmonella persisters undermine host immune defenses during antibiotic treatment. Science, 362(6419), 1156–1160. 10.1126/SCIENCE.AAT7148/SUPPL_FILE/AAT7148_TABLE_S9.XLSX

52. Stepien, T. A., Singletary, L. A., Guerra, F. E., Karlinsey, J. E., Libby, S. J., Jaslow, S. L., Gaggioli, M. R., Gibbs, K. D., Ko, D. C., Brehm, M. A., Greiner, D. L., Shultz, L. D., & Fang, F. C. (2024). Nuclear factor kappa B-dependent persistence of Salmonella Typhi and Paratyphi in human macrophages. MBio, 15(4), e00454–24. 10.1128/MBIO.00454-24

53. Subramanian, A., Tamayo, P., Mootha, V. K., Mukherjee, S., Ebert, B. L., Gillette, M. A., Paulovich, A., Pomeroy, S. L., Golub, T. R., Lander, E. S., & Mesirov, J. P. (2005). Gene set enrichment analysis: a knowledge-based approach for interpreting genome-wide expression profiles. Proceedings of the National Academy of Sciences of the United States of America, 102(43), 15545–15550. 10.1073/PNAS.0506580102

54. Tanigawa, Y., Dyer, E. S., & Bejerano, G. (2022). WhichTF is functionally important in your open chromatin data? PLoS Computational Biology, 18(8). 10.1371/JOURNAL.PCBI.1010378

55. Thurman, R. E., Rynes, E., Humbert, R., Vierstra, J., Maurano, M. T., Haugen, E., Sheffield, N. C., Stergachis, A. B., Wang, H., Vernot, B., Garg, K., John, S., Sandstrom, R., Bates, D., Boatman, L., Canfield, T. K., Diegel, M., Dunn, D., Ebersol, A. K., … Stamatoyannopoulos, J. A. (2012). The accessible chromatin landscape of the human genome. Nature 2012 489:7414, 489(7414), 75–82. 10.1038/nature11232

56. Weber-Nordtt, R. M., Riley, J. K., Greenlund, A. C., Moore, K. W., Darnell, J. E., & Schreiber, R. D. (1996). Stat3 Recruitment by Two Distinct Ligand-induced, Tyrosine-phosphorylated Docking Sites in the Interleukin-10 Receptor Intracellular Domain. Journal of Biological Chemistry, 271(44), 27954–27961. 10.1074/JBC.271.44.27954

57. Zaret, K. S., & Carroll, J. S. (2011). Pioneer transcription factors: establishing competence for gene expression. Genes & Development, 25(21), 2227–2241. 10.1101/GAD.176826.111

58. Zhong, Z., Wen, Z., & James E. Darnell, Jr. (1994). Stat3: a STAT Family Member Activated by Tyrosine Phosphorylation in Response to Epidermal Growth Factor and Interleukin-6. Science, 264(5155), 95–98. 10.1126/SCIENCE.8140422

