## Supplemental figures for "*Salmonella* Effector SteE Reprogrammes the Macrophage Regulatory Network to Drive Specific Hyperactivation of STAT3 Target Genes"

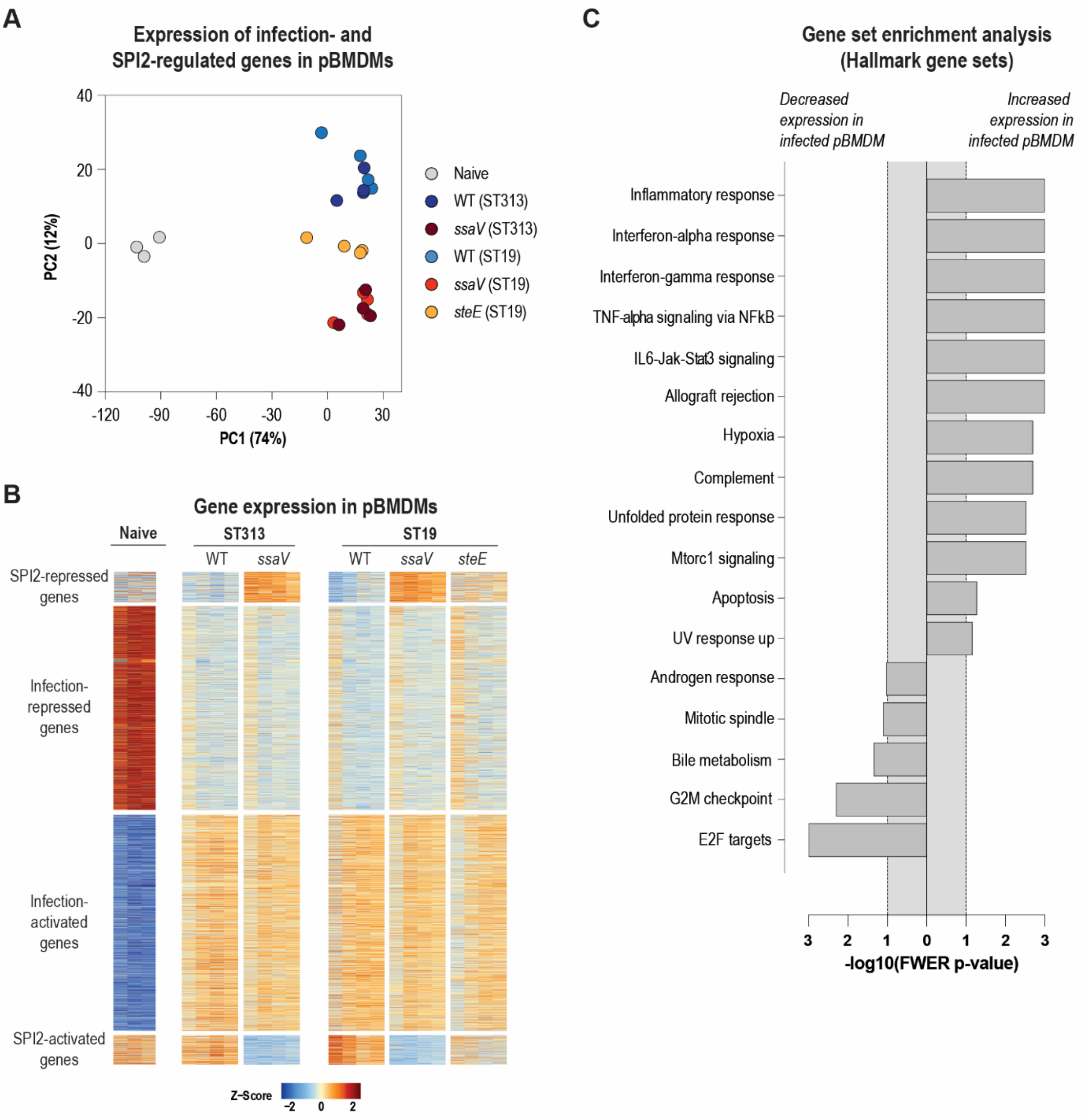


**Supplementary Figure 1 (related to Figure 1) – Gene expression dynamics in pBMDMs upon *Salmonella* Typhimurium infection.**

**(A-C)** Primary bone marrow-derived macrophages (pBMDMs) were untreated and unchallenged (naive), infected with wild-type or *ssaV*-mutant ST313 *Salmonella* Typhimurium (D23580), or infected with wild-type, *ssaV*-mutant or *steE*-mutant ST19 *Salmonella* Typhimurium (12023/14028). RNA sequencing was performed on samples taken 18 hours post-uptake.

**(A)** Principal component analysis of the transcriptomes of the indicated samples based on genes that are differentially expressed (adjusted *p*-value < 0.01, DESeq2) in pBMDMs infected with wild-type *Salmonella* Typhimurium compared to *ssaV* mutant-infected pBMDMs and in naïve pBMDMs compared with pBMDMs infected with wild-type *Salmonella* strains. The percentage of variance explained by the first principal component (PC1) and the second principal component (PC2) are shown.

**(B)** Heatmap showing clusters of genes that are differentially expressed (adjusted *p*-value < 0.01, DESeq2) in pBMDMs infected with wild-type *Salmonella* Typhimurium compared to *ssaV* mutant-infected pBMDMs and in naïve pBMDMs compared with pBMDMs infected with wild-type *Salmonella* strains. Genes are classified as SPI2-activated or SPI2-repressed and Infection-activated and Infection-repressed based on their expression patterns. Colours represent Z-scores, reflecting relative gene expression levels across all samples.

**(C)** Gene set enrichment analysis (GSEA) identifying hallmark pathways with significantly increased or decreased activity (FWER-adjusted *p*-value < 0.1). Comparisons were made between naïve pBMDMs and pBMDMs infected with wild-type *Salmonella* strains.

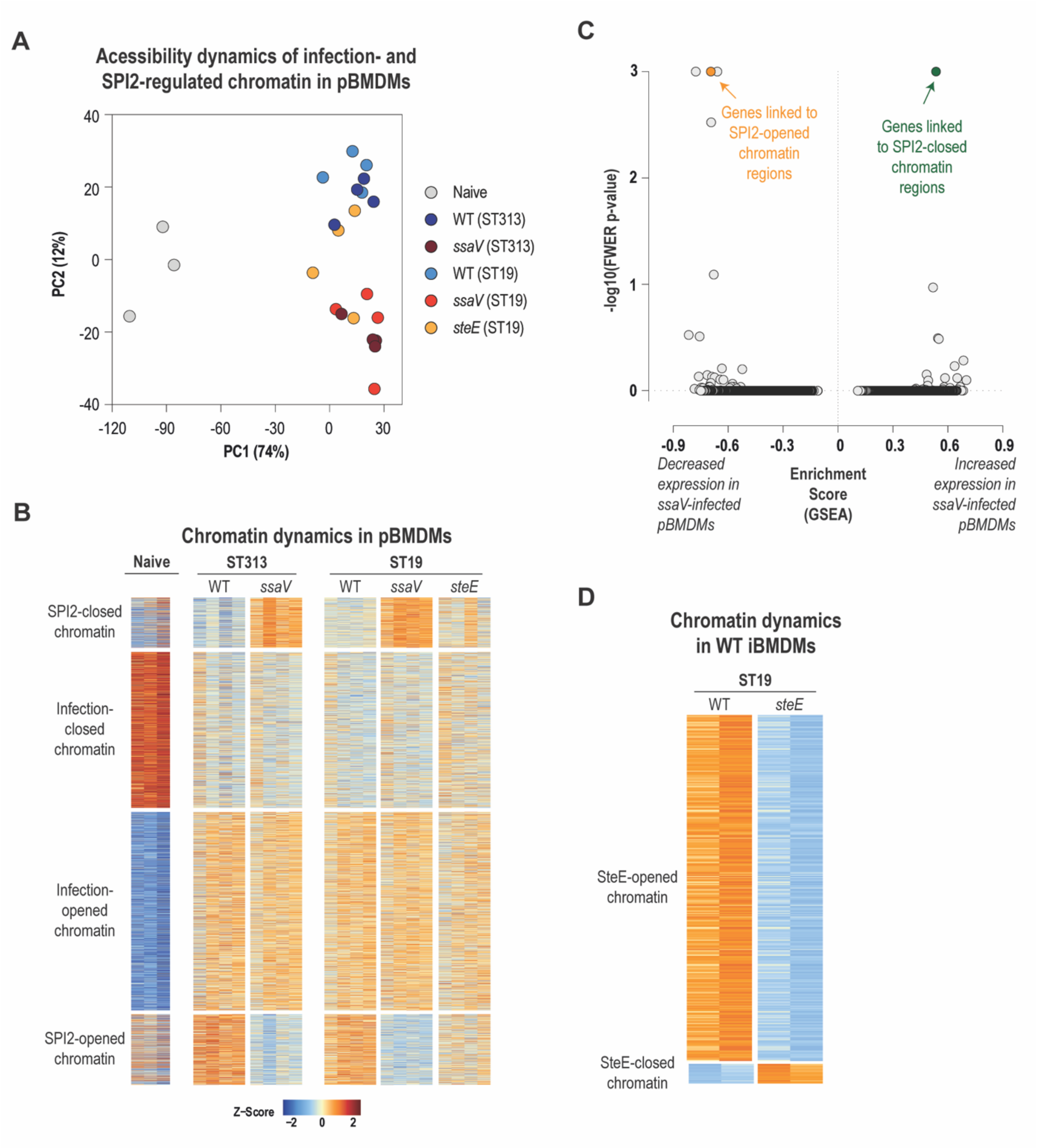


**Supplementary Figure 2 (related to Figure 2) – Chromatin accessibility in pBMDMs upon *Salmonella* Typhimurium infection.**

**(A-D)** Primary or immortalized bone marrow-derived macrophages (pBMDMs or iBMDMs) were untreated and unchallenged (naive), infected with wild-type or *ssaV*-mutant ST313 *Salmonella* Typhimurium (D23580), or infected with wild-type, *ssaV*-mutant or *steE*-mutant ST19 *Salmonella* Typhimurium (12023/14028). ATAC (**A**ssay for **T**ransposase **A**ccessible **C**hromatin) sequencing was performed on samples taken 18 hours post-uptake.

**(A)** Principal component analysis of chromatin regions that are differentially accessible (adjusted *p*-value < 0.1, DESeq2) in naïve pBMDMs compared with infected pBMDMs or in pBMDMs infected with wild-type *Salmonella* Typhimurium compared to *ssaV*-mutant infected pBMDMs (related to Figure 2A). The percentage of variance explained by the first principal component (PC1) and the second principal component (PC2) are shown.

**(B)** Heatmap showing clusters of chromatin regions that are differentially accessible (adjusted *p*-value < 0.1, DESeq2) in naïve pBMDMs compared with infected pBMDMs or in pBMDMs infected with wild-type *Salmonella* Typhimurium compared to *ssaV*-mutant infected pBMDMs. Regions are classified as SPI2-opened or SPI2-closed and Infection-opened or Infection-closed based on their accessibility patterns. Colours represent Z-scores, reflecting relative chromatin accessibility levels across all samples.

**(C)** Gene set enrichment analysis (GSEA) identifying macrophage gene sets with increased or decreased expression (based on RNA-Seq; see Figure 1) in pBMDMs infected with *ssaV*-mutant *Salmonella* Typhimurium strains compared to those infected with the wild-type strains. Gene sets linked to SPI2-opened and SPI2-closed chromatin regions (see Figure 2A) were identified using the Genomic Regions Enrichment of Annotations Tool (GREAT). Additional gene sets, including all hallmark gene sets, canonical pathway gene sets, and gene ontology (GO) gene sets, were sourced from GSEA. The plot shows the GSEA enrichment score (x-axis) and the -log10(FWER-adjusted *p*-value) (y-axis) for all gene sets analysed.

**(D)** Heatmap showing clusters of chromatin regions that are differentially accessible (adjusted *p*-value < 0.1, limma) in wild-type iBMDMs infected with wild-type ST19 *Salmonella* Typhimurium compared to those infected with an isogenic *steE*-mutant strain. Genes are classified as SteE-opened or SteE-closed based on their accessibility patterns. Colours represent Z-scores, reflecting relative chromatin accessibility levels across all samples.

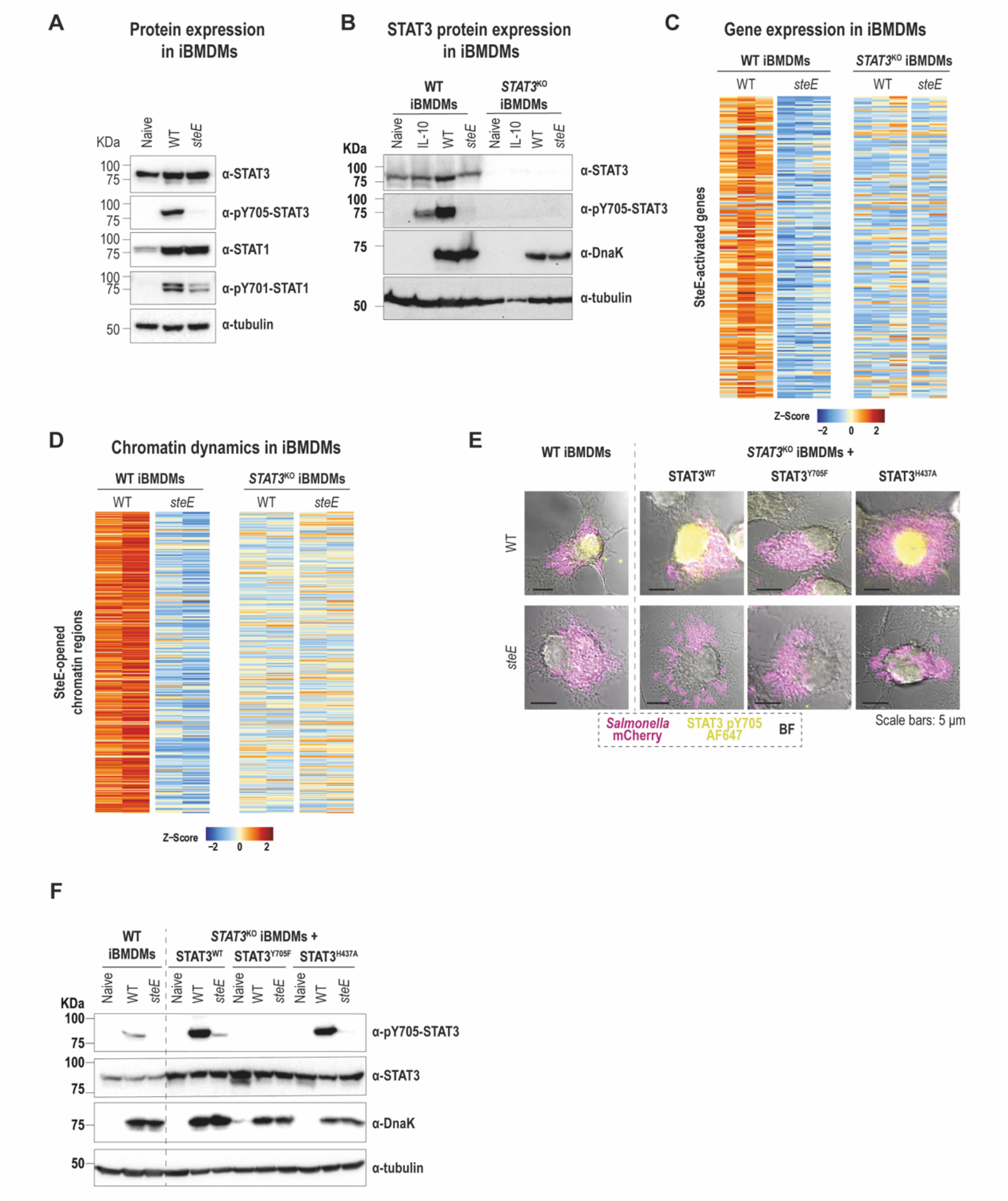


**Supplementary Figure 3 (related to Figure 3) – SteE-induced gene expression and chromatin accessibility patterns, and STAT3 protein levels and localisation, in wild type and *STAT3*$^{KO}$ iBMDMs.**

**(A-F)** Wild type (WT) and *STAT3*$^{KO}$ immortalized bone marrow-derived macrophages (iBMDMs) were untreated and unchallenged (naive), infected with wild-type or *steE*-mutant ST19 *Salmonella* Typhimurium strains, or treated with 20 ng/ml IL10. Assays were performed 18 hours post-uptake or post-treatment.

**(A)** Immunoblot analysis of indicated proteins in iBMDM. Cell extracts were examined for STAT3 and STAT1 protein expression and tyrosine phosphorylation, with tubulin, used as control. Data shown are representative of three independent experiments.

**(B)** Immunoblot analysis of STAT3 in macrophages naïve, infected and IL10-treated wild type iBMDMs (left) and *STAT3*$^{KO}$ iBMDMs (right). Cell extracts were examined for STAT3 protein expression and tyrosine phosphorylation, with tubulin and DnaK used as controls. Data shown are representative of three independent experiments.

**(C)** Heatmap showing genes differentially activated (adjusted p-value < 0.01, DESeq2) in iBMDMs infected with wild-type ST19 *Salmonella* Typhimurium compared to iBMDMs infected with an isogenic *steE*-mutant strain. Results are shown for wild type iBMDMs (left) and *STAT3*$^{KO}$ iBMDMs (right). Colours represent Z-scores, reflecting relative gene expression across all samples.

**(D)** Heatmap showing chromatin regions that are differentially opened (adjusted $p$-value < 0.1, limma) in iBMDMs infected with wild-type *Salmonella* Typhimurium compared to iBMDMs infected with an isogenic *steE*-mutant strain. Results are shown for wild type iBMDMs (left) and *STAT3*$^{KO}$ iBMDMs (right). Colours represent Z-scores, reflecting relative chromatin accessibility levels across all samples.

**(E)** Immunofluorescence microscopy images of wild type or *STAT3*$^{KO}$ iBMDMs complemented with STAT3$^{WT}$, STAT3$^{Y705F}$ or STAT3$^{H437A}$, infected with wild-type or *steE* ST19 *Salmonella* Typhimurium at 18 h post-uptake. Merged images represent *Salmonella* (mCherry, Magenta) pY705-STAT3 (Yellow, AF647) and brightfield (grey). Scale bars are 5 µm. Data shown is representative of two independent experiments.

**(F)** Immunoblot analysis of STAT3 protein expression. Wild type (WT) or *STAT3*$^{KO}$ iBMDMs complemented with STAT3$^{WT}$, STAT3$^{Y705F}$ or STAT3$^{H437A}$ were either untreated and unchallenged (naïve) or infected with wild-type or *steE*-mutant ST19 *Salmonella* Typhimurium. Cell lysates were examined for STAT3 protein expression and tyrosine phosphorylation, with DnaK, and tubulin as controls. Data shown is representative of two independent experiments.

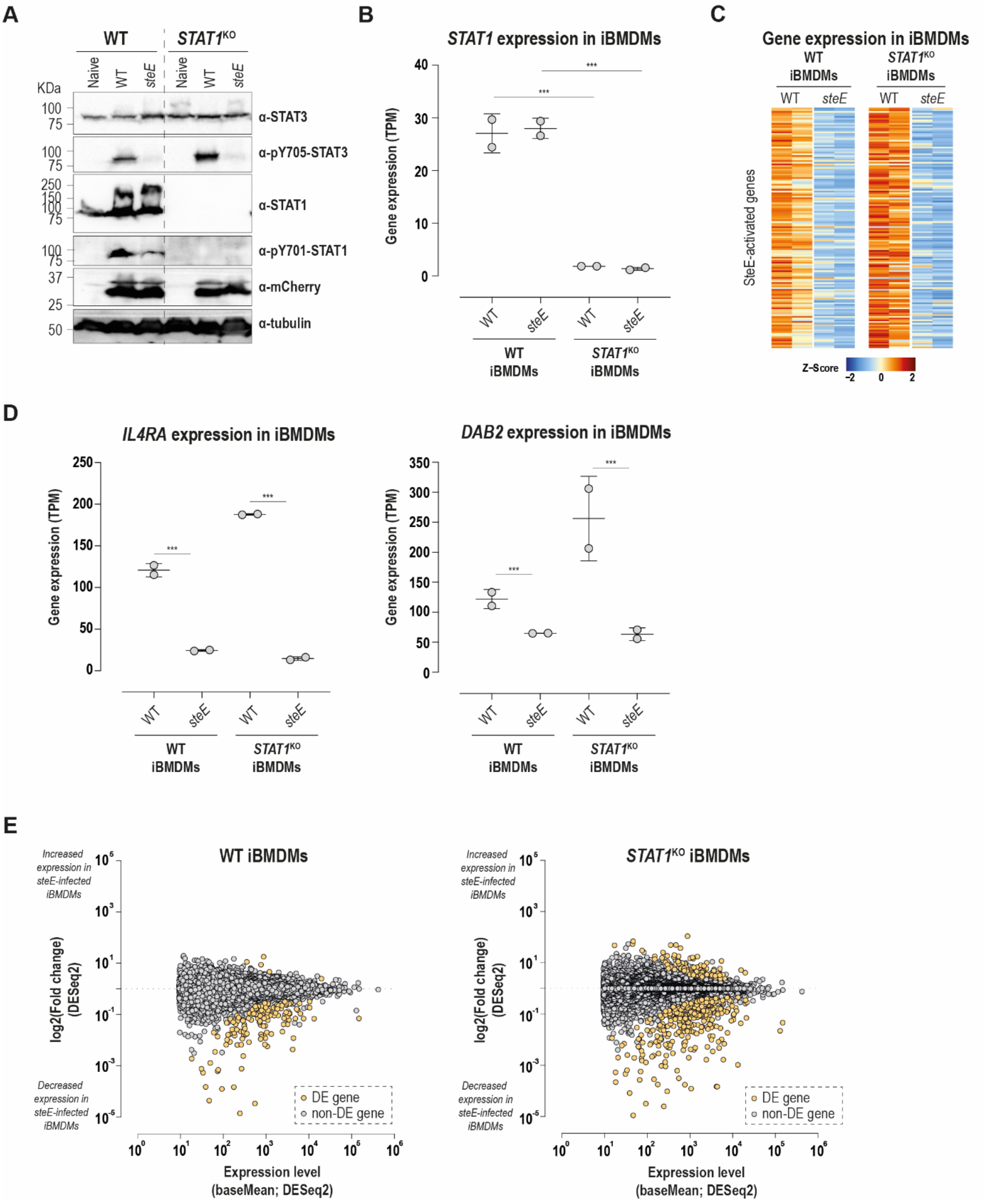


**Supplementary Figure 4 (related to Figure 3) – STAT1 expression and SteE-induced gene expression patterns in wild type and *STAT1*$^{KO}$ iBMDMs.**

**(A-E)** Wild-type (WT) and *STAT1*$^{KO}$ immortalized bone marrow-derived macrophages (iBMDMs) were untreated and unchallenged (naïve) or infected with wild-type or *steE*-mutant ST19 *Salmonella* Typhimurium strains. Assays were performed on samples taken 18 hours post-uptake.

**(A)** Immunoblot analysis of STAT1 and STAT3 protein expression in wild-type (WT) or *STAT1*$^{KO}$ iBMDMs. Cell lysates were examined by immunoblot for STAT1 and STAT3 protein expression and tyrosine phosphorylation, with mCherry, and tubulin as controls. Data shown is representative of two independent experiments.

**(B)** Gene expression levels for STAT1 in wild type and *STAT1*$^{KO}$ iBMDMs. Differential gene expression was analysed using DESeq2, with adjusted p-values shown (∗∗∗ adj. $p < 0.001$).

**(C)** Heatmap visualising gene expression levels in wild type or *STAT1*$^{KO}$ iBMDMs harbouring either wild-type or *steE* ST19 *Salmonella* Typhimurium at 18 h post-uptake for SteE-activated genes in iBMDMs.

**(D)** Gene expression levels for the anti-inflammatory marker genes *IL4RA* (left) and *DAB2* (right), in wild type and *STAT1*$^{KO}$ iBMDMs infected with wild-type ST19 *Salmonella* Typhimurium or an isogenic *steE*-mutant strain. RNASeq was performed 18 hours post-uptake. Differential gene expression was analysed using DESeq2, with adjusted p-values shown (∗∗∗ adj. $p < 0.001$).

**(E)** Scatterplots illustrating differential gene expression between iBMDMs infected with wild-type ST19 *Salmonella* Typhimurium and an isogenic *steE*-mutant strain. Data are shown for wild type (left panel), and *STAT1*$^{KO}$ iBMDMs (right panel). The shrunken $\log_2$ fold change (y-axis, DESeq2), and the mean expression level (x-axis, DESeq2) are shown. Yellow dots highlight differentially expressed genes (adjusted p-value < 0.01, DESeq2).

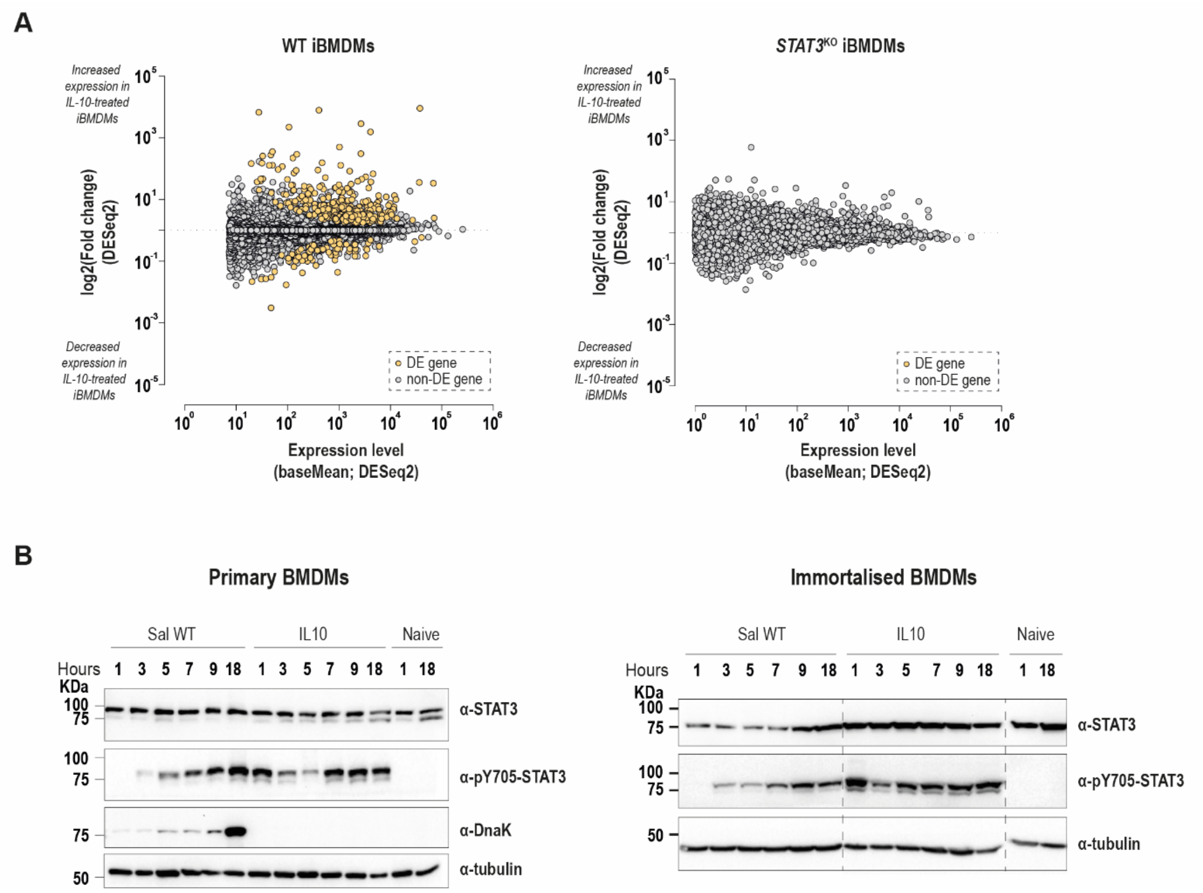


**Supplementary Figure 5 (related to Figure 4) – Gene expression patterns in *STAT3*$^{KO}$ iBMDMs upon IL10 treatment, and population level STAT3 phosphorylation levels in infected or IL10-treated iBMDMs**

**(A)** Scatterplots illustrating differential gene expression between naïve (untreated and unchallenged) and IL10-treated macrophages. Data are shown for wild type iBMDMs (left panel) and *STAT3*$^{KO}$ iBMDMs (right panel). The shrunken $\log_2$ fold change (y-axis, DESeq2), and the mean expression level (x-axis, DESeq2) are shown. Yellow dots highlight differentially expressed genes (adjusted p-value < 0.01, DESeq2).

**(B)** Immunoblot analysis of STAT3 phosphorylation in macrophages. pBMDMs (left) or iBMDMs (right) were either naïve (untreated and unchallenged), infected with wild-type ST19 *Salmonella* Typhimurium, or treated with 20 ng/mL IL10 for the indicated time. Cell lysates were examined by immunoblotting for STAT3 protein expression and tyrosine phosphorylation, with tubulin and DnaK used as controls. Data shown are representative of two independent experiments. Dotted lines on the right panel represent a post analysis change in lane order, with all samples shown from one continuous membrane.
